# Biological Sequence Modeling with Convolutional Kernel Networks

**DOI:** 10.1101/217257

**Authors:** Dexiong Chen, Laurent Jacob, Julien Mairal

## Abstract

The growing number of annotated biological sequences available makes it possible to learn genotype-phenotype relationships from data with increasingly high accuracy. When large quantities of labeled samples are available for training a model, convolutional neural networks can be used to predict the phenotype of unannotated sequences with good accuracy. Unfortunately, their performance with medium- or small-scale datasets is mitigated, which requires inventing new data-efficient approaches. In this paper, we introduce a hybrid approach between convolutional neural networks and kernel methods to model biological sequences. Our method enjoys the ability of convolutional neural networks to learn data representations that are adapted to a specific task, while the kernel point of view yields algorithms that perform significantly better when the amount of training data is small. We illustrate these advantages for transcription factor binding prediction and protein homology detection, and we demonstrate that our model is also simple to interpret, which is crucial for discovering predictive motifs in sequences. The source code is freely available at https://gitlab.inria.fr/dchen/CKN-seq.

## 1 Introduction

Understanding the relationship between biological sequences and the associated phenotypes is a fundamental problem in molecular biology. Accordingly, machine learning techniques have been developed to exploit the growing number of phenotypic sequences in automatic annotation tools. Typical applications include classifying protein domains into superfamilies [Leslie et al., 2003, Saigo et al., 2004], predicting whether a DNA or RNA sequence binds to a protein [Alipanahi et al., 2015], its splicing outcome [Jha et al., 2017], or its chromatin accessibility [Kelley et al., 2016], predicting the resistance of a bacterial strain to a drug [Drouin et al., 2016], or denoising a ChIP-seq signal [Koh et al., 2017].

Choosing how to represent biological sequences is a critical part of methods that predict phenotypes from genotypes. Kernel-based methods [Schölkopf and Smola, 2002] have often been used for this task. Biological sequences are represented by a large set of descriptors, constructed for instance by Fisher score [Jaakkola et al., 2000], k-mer spectrum up to some mismatches [Leslie et al., 2003], or local alignment score [Saigo et al., 2004]. By using the so-called kernel trick, these huge-dimensional descriptors never need to be explicitly computed as long as the inner-products between pairs of such vectors can be efficiently computed. A major limitation of traditional kernel methods is their use of fixed representations of data, as opposed to optimizing representations for a specific task. Another issue is their poor scalability since they require computing a *n* × *n* Gram matrix where *n* is the number of data points.

By contrast, methods based on convolutional neural networks (CNN) are more scalable and are able to optimize data representations for a specific prediction problem [LeCun et al., 1989]. Even though their predictive performance was first demonstrated for two-dimensional images, they have been recently successfully adopted for DNA sequence modeling [Alipanahi et al., 2015, Zhou and Troyanskaya, 2015]. When sufficient annotated data is available, they can lead to good prediction accuracy, though they still suffer from some known limitations. An important one is their lack of interpretability: the set of functions described by the network is only characterized by its algorithmic construction, which makes both the subsequent analysis and interpretation difficult. CNNs for DNA sequences typically involve much fewer layers than CNNs for images, and lend themselves to some level of interpretation [Alipanahi et al., 2015, Lanchantin et al., 2017, Shrikumar et al., 2017a]. However, a systematic approach is still lacking as existing methods rely on specific sequences to interpret trained filters [Alipanahi et al., 2015, Shrikumar et al., 2017a] or output a single feature per class [Lanchantin et al., 2017, (3.3)]. Correctly regularizing neural networks to avoid overfitting is another open issue and involves various heuristics such as dropout [Srivastava et al., 2014], weight decay [Hanson and Pratt, 1989], and early stopping. Finally, training neural networks generally requires large amounts of labeled data. When few training samples are available, training CNNs is challenging, motivating us for proposing a more data-efficient approach.

In this paper we introduce CKN-seq, a strategy combining kernel methods and deep neural networks for sequence modeling, by adapting the convolutional kernel network (CKN) model originally developed for image data [Mairal, 2016]. CKN-seq relies on a continuous relaxation of the mismatch kernel [Leslie and Kuang, 2004]. The relaxation makes it possible to learn the kernel from data, and we provide an unsupervised and a supervised algorithm to do so – the latter being a special case of CNNs. On the datasets we consider, both approaches show better performance than DeepBind, another existing CNN [Alipanahi et al., 2015], especially when the amount of training data is small. On the other hand, the supervised algorithm produces task-specific and small-dimensional sequence representations while the unsupervised version dominates all other methods on small-scale problems but leads to higher dimensional representations. Consequently, we introduce a *hybrid* approach which enjoys the benefits of both supervised and unsupervised variants, namely the ability of learning low-dimensional models with good prediction performance in all data size regimes. Finally, the kernel point of view of our method provides us simple ways to visualize and interpret our models, and obtain sequence logos.

We investigate the performance of CKN-seq on a transcription factor binding prediction task as well as on a protein remote homology detection. We provide a free implementation of CKN-seq for learning from biological sequences, which can easily be adapted to other sequence prediction tasks.

## 2 Method

In this section, we introduce our approach to learning sequence representations. We first review CNNs and kernel methods over which our convolutional kernel network is built. Then, we present the construction of CKN followed by the learning method. We finish the section with discussions on the interpretation and visualization of a trained CKN.

### 2.1 Supervised learning problem

Let us consider *n* sequence samples **x**_1_, **x**_2_, … , **x**_*n*_ in a set 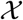 of variable-length biological sequences. The sequences are assumed to be over an alphabet 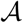. Each sequence **x**_*i*_ is associated to a measurement *y*_*i*_ in 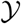 denoting some biological property of the sequence. For instance, 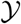 may be binary labels {−1, 1} (*e.g.*, whether the sequence is bound by a particular transcription factor or not) or ℝ for continuous traits (*e.g.*, the expression of a gene). The goal of supervised learning is to use these *n* examples {**x**_*i*_, *y*_*i*_}_*i*=1,…,*n*_ to learn a function 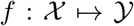 which accurately predicts the label of a new, unobserved sequence. Learning is typically achieved by minimizing the following objective:

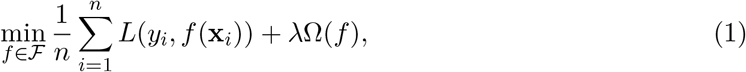

where *L* is a loss function measuring how well the prediction *f*(**x**_*i*_) fits the true label *y*_*i*_, and Ω measures the smoothness of *f*. 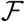 is a set of candidate functions over which the optimization is performed. Both CNNs and kernel methods can be thought of as manners to design this set.

#### Convolutional neural networks

In neural networks, the functions in 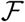 perform a sequence of linear and nonlinear operations that are interleaved in a multilayer fashion. Specifically, the CNN DeepBind [Alipanahi et al., 2015] represents the four DNA characters respectively as the vectors (1, 0, 0, 0), (0, 1, 0, 0), (0, 0, 1, 0), (0, 0, 0, 1), such that an input sequence **x** of length *m* is represented as a 4×*m* matrix. DeepBind then produces an intermediate representation obtained by one-dimensional convolution of the full sequence **x** with *p* convolution filters, followed by a pointwise non-linear function and a max pooling operation along each sequence, yielding a representation **x̃** in ℝ^*p*^ of the sequence. A final linear prediction layer is applied to **x̃**. The optimization in (1) acts on both the weights of this linear function and the convolution filters. Therefore, DeepBind simultaneously learns a representation **x̃** and a linear prediction function over this representation.

DeepBind additionally modifies the objective function (1) to enforce an invariance to reverse complementation of **x**. The loss term is replaced with *L* (*y*_*i*_, max (*f*(**x**_*i*_, f(**x̄**_*i*_))) where **x̄** denotes the reverse complement of **x**. Using this formulation is reported by Alipanahi et al. [2015] to improve the prediction performance. Other versions have been then considered, by using a fully connected layer that allows mixing information from the two DNA strands [Shrikumar et al., 2017b], or by considering several hidden layers instead of a single one [Zeng et al., 2016]. Overall, across several versions, the performance of DeepBind with a single hidden layer turned out to be the best on average on ChIP-seq experiments from ENCODE [Zeng et al., 2016].

#### Kernel methods

Like in CNNs, the main principle of kernel methods is to implicitly map each training point **x**_*i*_ to a feature space in which simpler predictive functions are applied. For kernel methods, these feature spaces are generally high- (or even infinite-) dimensional vector spaces. This is achieved indirectly, by defining a kernel function 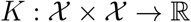 which acts as a similarity measure between input data. When the kernel function is symmetric and positive definite, a classical result [see Schölkopf and Smola, 2002] states that there exists a Hilbert space 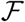 of functions from 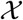 to ℝ, called reproducing kernel Hilbert space (RKHS), along with a mapping 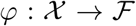, such that 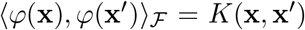 for all (**x**, **x**′) in 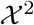, where 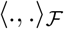 is the Hilbertian inner-product associated with 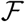. In other words, there exists a mapping of sequences into a Hilbert space, such that the kernel value between any sequence pairs is equal to the inner-product between their maps in the Hilbert space. Besides, any function *f* in 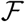 may be interpreted as a linear form 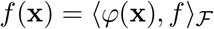 for all **x** in 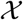. A large number of kernels have been specifically designed for biological sequences [see Ben-Hur et al., 2008, and references therein].

In the context of supervised learning (1), training points **x**_*i*_ can be mapped into *φ*(**x**_*i*_) in 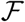, and we look for a prediction function *f* in 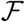. Interestingly, regularization is also convenient in the context of kernel methods, which is crucial for learning when few labeled samples are available. By choosing the regularization function 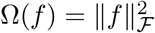, it is indeed possible to control the regularity of the prediction function *f*: for any two points **x**, **x′** in 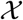, the variations of the predictions are bounded by 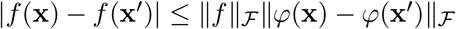. Hence, a small norm 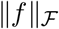 implies that *f*(**x**) will be close to *f*(**x′**) whenever **x**and **x′**are close to each other according to the geometry induced by the kernel.

Kernel methods have several assets: (i) they are generic and can be directly applied to any type of data – *e.g.*, sequences or graphs – as long as a relevant positive definite kernel is available; (ii) they are easy to regularize. However, as alluded earlier, naive implementations lack scalability. A typical workaround is the Nyström approximation [Williams and Seeger, 2001], which builds an explicit *q*-dimensional mapping 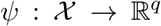 for a reasonably small *q* approximating the kernel, *i.e.*, such that 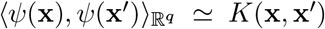. Then, solving the regularized problem (1) under this approximation amounts to learning a linear model with *q* dimensions. We will discuss how CKNs circumvent the scalability problem, while being capable to produce task-adapted data representations.

### 2.2 Convolutional kernel networks for sequences

We introduce convolutional kernel networks for sequences, and show their link with mismatch kernels [Leslie and Kuang, 2004].

#### 2.2.1 Convolutional kernel for sequences

Given two sequences **x** and **x′** of respective lengths *m* and *m*′, we consider a window size *k*, and we define the following kernel, which compares pairwise subsequences of length *k* (*k*-mers) within **x** and **x′**:

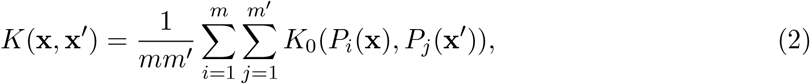

where *P*_*i*_(**x**) is a *k*-mer of **x** centered at position *i*, represented as a one-hot encoded vector of size 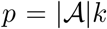 and *K*_0_ is a positive definite kernel used to compare *k*-mers.^1^ We follow Mairal [2016] and use a homogeneous dot-product kernel such that for two vectors **z** and **z′** in ℝ^*p*^,

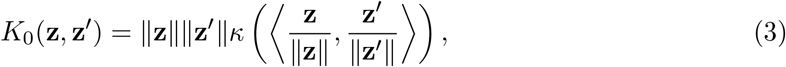

and 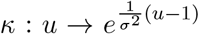. Note that when **z** and **z′** are one-hot encoded vectors of subsequences, 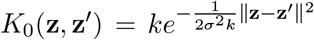 (more details can be found in Supplementary Section A), and we recover a Gaussian kernel that involves the Hamming distance ‖**z** − **z′**‖^2^/2 between the two sub-sequences. Up to the normalization factors, this choice leads to the same kernel used by Morrow et al. [2017]. Yet, the algorithms we will present next are significantly different. While Morrow et al. [2017] use random features [Rahimi and Recht, 2008] to find a finite-dimensional mapping 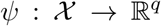 that approximates the kernel map, our approach relies on the Nyström approximation [Williams and Seeger, 2001]. A major advantage of the Nyström method is that it may be extended to produce lower-dimensional *task-dependent* mappings [Mairal, 2016] and it admits a model interpretation in terms of sequence logos (see Section 3).

#### 2.2.2 Learning sequence representation

The positive definite kernel *K*_0_ defined in (3) implicitly defines a reproducing kernel Hilbert space 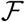 over *k*-mers, along with a mapping 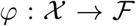. The convolutional kernel network model uses the Nyström method to approximate any point in 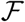 onto its projection on a finite-dimensional subspace *ε* defined as the span of some anchor points

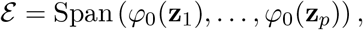

where the **z**_*i*_’s are the anchor points in 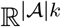. Subsequently, it is possible to define a coordinate system in *ε* such that the orthogonal projection of *φ*_0_(**z**) onto *ε* may be represented by a *p*-dimensional vector *ψ*_0_(**z**). Assume for now that the anchor points **z**_*i*_ are given. Then, a finite-dimensional embedding [see Mairal, 2016, for details] is given by

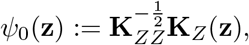

where 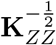 is the inverse (or pseudo inverse) square root of the *p* × *p* Gram matrix [*K*_0_(**z**_*i*_, **z**_*j*_)]_*ij*_ and **K**_*Z*_(**z**) = (*K*_0_(**z**_1_, **z**),…, *K*_0_(**z**_*p*_, **z**))^⊤^. It is indeed possible to show that this vector preserves the Hilbertian inner-product in 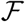 after projection: 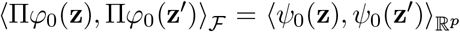 for any **z**, **z′** in 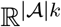, where Π denotes the orthogonal projection onto *ε*. Assuming *P*_*i*_(**x**) and *P*_*j*_(**x′**) map close enough to *ε*, a reasonable approximation is therefore 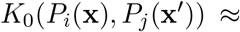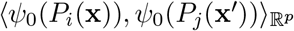 for all *i*, *j* in (2), and then

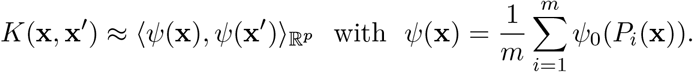

Finally, the original optimization problem (1) can be approximated by

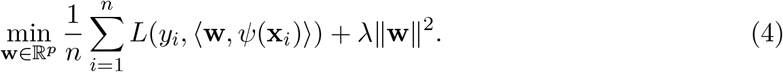

We have assumed so far that the anchor points **z**_*i*_, *i* = 1 …, *p* were given – *i.e.*, that the sequence representation *ψ*(**x**) was fixed in advance. We now present two methods to learn this representation. The overall approximation scheme is illustrated in the left panel of Figure 1.

**Figure 1.**
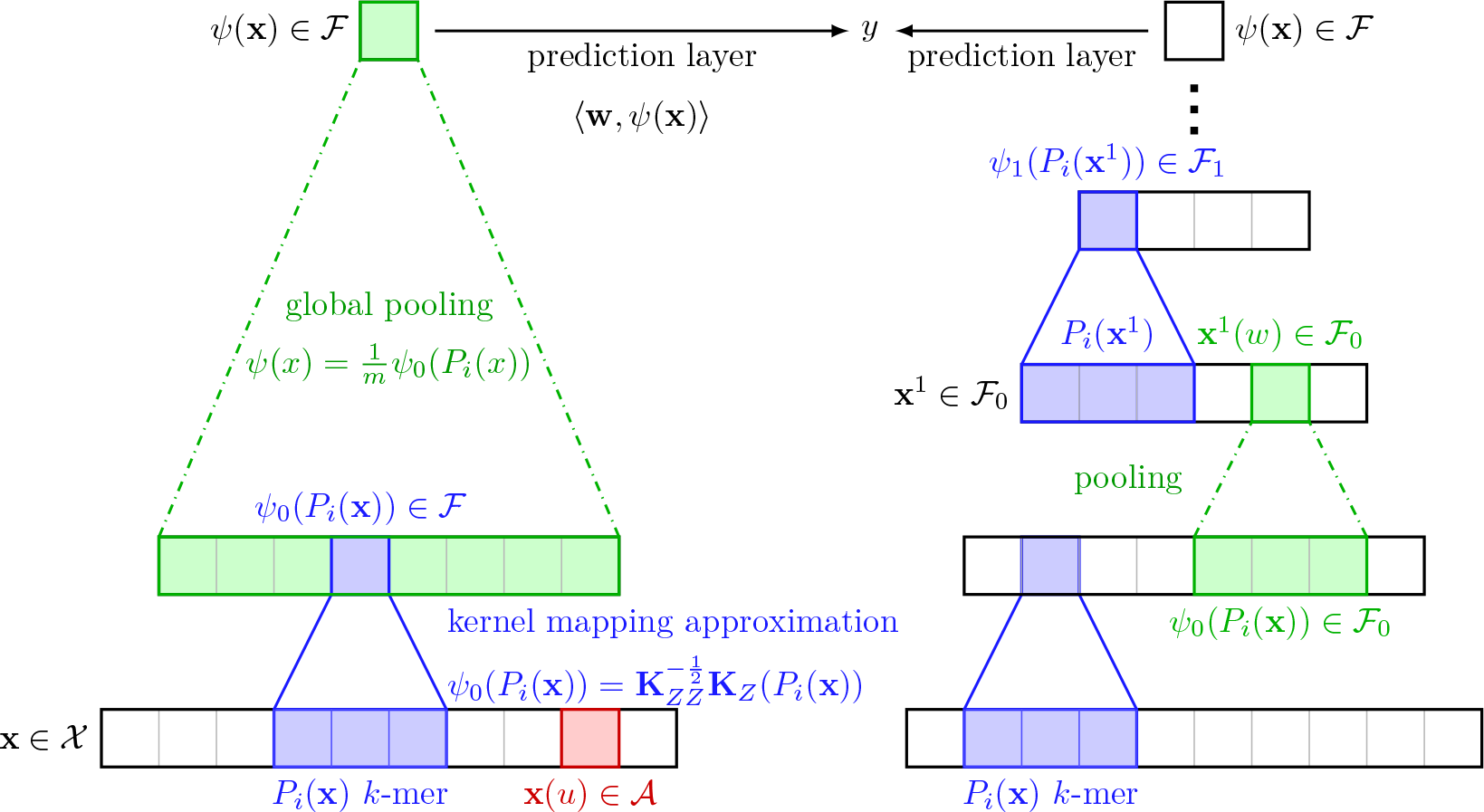
Construction of single-layer (left) and multilayer (right) CKN-seq. For a single-layer model, each *k*-mer *P*_*i*_(**x**) is mapped to *φ*_0_(*P*_*i*_(**x**)) in 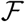 and projected to 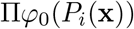 parametrized by *ψ*_0_(*P*_*i*_(**x**)). Then, the final finite-dimensional sequence is obtained by the global pooling, 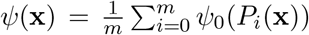. The multilayer construction is similar, but relies on intermediate maps, obtained by local pooling, see main text for details.

##### Unsupervised learning of the anchor points

The first strategy consists in running a clustering algorithm such as K-means in order to find *p* centroids **z**_*i*_ in 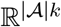 that “span” well the data. This is achieved by extracting a large number of *k*-mers from the training sequences and by clustering them. The method is simple, performs well in practice as shown in Section 3, and can also be used to initialize the training of the following supervised variant. However, the main drawback is that it generally requires a large number of anchor points (see Section 3) to achieve good prediction, which can be problematic for model interpretation.

##### Supervised learning of the anchor points

The other strategy consists in jointly optimizing (4) with respect to the vector **w** in ℝ^*p*^ and to the anchor points that parametrize the representation *ψ*.

In practice, we adopt an optimization scheme that alternates between two steps: (a) we fix the anchor points (**z**_*i*_)_*i*=1,=,*p*_, compute the finite-dimensional representations *ψ*(**x**_1_), … , *ψ*(**x**_*n*_) of all data points, and minimize function (4) with respect to **w**, which is convex if *L* is convex; (b) We fix **w** and update all the (**z**_*i*_)_*i*=1,…,*p*_ using one pass of a projected stochastic gradient descent (SGD) algorithm while fixing **w**, at a similar computational cost per iteration as a classical CNN. The optimization for the reverse-complement formulation can be done in the same way except that it is no more convex with respect to **w**, but we can still apply a fast optimization method such as L-BFGS [Liu and Nocedal, 1989]. We find this alternating scheme more efficient and more stable than using an SGD algorithm jointly on **w**and the anchor points.

#### 2.2.3 Multilayer construction

We have presented CKNs with a single layer for simplicity, but the extension to multiple layers is straightforward. Instead of reducing the intermediate representation in the left panel of Figure 1 to a single point, the pooling operation may simply reduce the sequence length by a constant factor (right panel of Figure 1), in a similar way as pooling reduces image resolution in CNN. This leads to an intermediate sequence representation **x**^1^ and we can define a valid kernel *K*_1_, the same as *K*_0_ in (3), but on subsequences of **x**^1^. Then the same approximation described in Section 2.2.2 can be applied to *K*_1_. In this way, the previous process can be repeated and stacked several times, by defining a sequence of kernels *K*_1_*, K*_2_, … on subsequences from the previous respective layer representations, along with Hilbert spaces 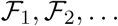 and mapping functions *φ*_1_, *φ*_2_,… [see Mairal, 2016]. Going up in the hierarchy, each point would carries information from a larger sequence neighborhood with more invariance due to the effect of pooling layers [Bietti and Mairal, 2017]. The training strategy is the same as for single-layer models.

Multilayer networks can potentially model larger motifs, with larger receptive fields, and possibly discover more interesting nonlinear relations between input variables than single-layer models. However, for the transcription factor binding prediction task under the setting of DeepBind or Zeng et al. [2016], we have observed that increasing the number of convolutional layers for CKN-seq did not improve the predictive performance (Supplementary Figure 8), as also observed by Zeng et al. [2016] for CNNs. The use of multiple layers may be however important when processing very long sequences, as observed for instance by Kelley et al. [2018], who also use dilated convolutions to model even larger receptive fields than what regular CNNs can achieve.

#### 2.2.4 Difference between supervised CKNs and CNNs

The main differences between CKN and CNN models are the choice of activation function (we used an exponential function in our experiments: *κ*(*x*) = *e*^*α*(*x*−1)^) and the transformation by the inverse square root of the Gram matrix. From a kernel point of view, the inverse square root of the Gram matrix allows us to interpret the operation as a projection onto a finite-dimensional subspace of an RKHS. From a neural network point of view, this operation decorrelates the channel entries. This can be observed when using a linear activation function *κ*(*u*) = *u*. In such a case, the approximated mapping is then 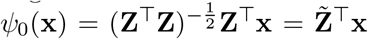, where 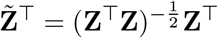 is an orthogonal matrix. Encouraging orthogonality of the filters has been shown useful to regularize deep networks [Cisse et al., 2017], and may provide intuition why our models perform better when small amounts of labeled data are available.

### 2.3 Data-augmented and hybrid CKN

As shown in our experiments, the unsupervised variant is sometimes more effective than the supervised one when there are only few training samples. In this section, we present a hybrid approach that can achieve similar performance as the unsupervised variant, while keeping a low-dimensional sequence representation that is easier to interpret. Before introducing this approach, we first present a classical data augmentation method for sequences, which consists in artificially generating additional training data, by perturbing the existing training samples. Formally, we consider random perturbations *δ*, such that given a sequence represented by a one-hot encoded vector **x**, we denote by **x** + *δ* the one-hot encoding vector of a perturbed sequence obtained by randomly changing some characters. Each character is switched to a different one, randomly chosen from the alphabet, with some probability *p*. With such a data augmentation strategy, the objective (1) then becomes

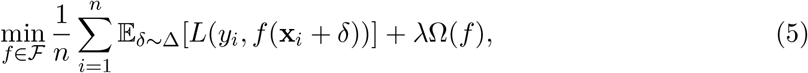

where Δ is a probability distribution of the variables *δ* corresponding to the perturbation process described above. The main assumption is that a perturbed sequence **x**_*i*_ + *δ* should have the same phenotype *y*_*i*_ when the perturbation *δ* is small enough. Whereas such an assumption may not be justified in general from a biological point of view, it led to significant improvements in terms of predictive accuracy. One possible explanation may be that for the tasks we consider, determining sequences may be short compared to the entire sequence: changing a few uniformly sample positions is therefore unlikely to perturb key bases.

As we show in Section 3, data-augmented CKN performs significantly better than its unaugmented counterpart when the amount of data is small. Yet, the unsupervised variant of CKN appears to be easier to regularize, and sometimes outperform all other approaches in such a low-data regime. This observation motivates us to introduce the following hybrid variant. In a first step, we learn a prediction function *f*_*u*_ based on the unsupervised variant of CKN, which leads to a high-dimensional sequence representation with good predictive performance. Then, we learn a low-dimensional model *f*_*s*_, whose purpose is to mimic the prediction of *f*_*u*_, by minimizing the cost function

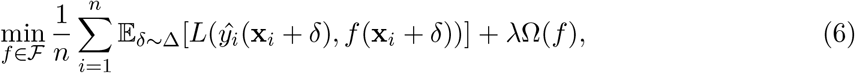

where 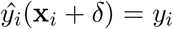 if *δ* = 0 and *f*_*u*_(**x**_*i*_ + *δ*) otherwise. Typically, the amount of perturbation that formulation (6) can afford is much larger than (5), as shown in our experiments, since it does not require to make the assumption that the sequence **x**_*i*_ + *δ* should have exactly label *y*_*i*_, which is a wrong assumption when *δ* is large.

### 2.4 Model interpretation and visualization

As observed by Morrow et al. [2017], the mismatch kernel [Leslie and Kuang, 2004] for modeling sequences may be written as Eq. (2) when replacing *K*_0_ with a discrete function *I*_0_ that assesses whether the two *k*-mers are identical up to some mismatches. Thus, the convolutional kernel (2) can be viewed as a continuous relaxation of the mismatch kernel. Such a relaxation allows us to characterize the approximated convolutional kernel by the learned anchor points (the variables **z**_1_, … , **z**_*p*_ in Section 2.2.2) that can be written as matrices in 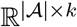.

To transform these optimized anchor points **z**_*i*_ into position weight matrices (PWMs) which can then be visualized as sequence logos, we identify the closest PWM to each **z**_*i*_: the kernel *K*_0_ implicitly defines a distance between one-hot-encoded sequences of length *k*, which is approximated by the Euclidean norm after mapping with *ψ*_0_. Given an anchor point **z**_*i*_, the closest PWM **μ** according to the geometry induced by the kernel is therefore obtained by solving

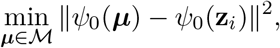

where 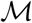 is the set of matrices in 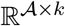 whose columns sum to one. This projection problem can be solved using a projected gradient descent algorithm. The simplicial constraints induce some sparsity to the resulting PWM, yielding more informative logos. As opposed to the approach of Alipanahi et al. [2015] which has relied on extracting *k*-mers sufficiently close to the filters in a validation set of sequences, the results obtained by our method do not depend on a particular dataset.

## 3 Application

We now study the effectiveness of CKN-seq on a transcription factor (TF) binding prediction and a protein homology detection problem.

### 3.1 Prediction of transcription factor binding sites

The problem of predicting TF binding sites has been extensively studied in the recent years with the continuously growing number of TF-binding datasets. This problem can be modeled as a classification task where the input is some short DNA sequence, and the label indicates whether the sequence can be bound by a TF of interest. It has recently been pointed out that incorporating non-sequence-based data modalities such as chromatin state can improve TF binding prediction [Karimzadeh and Hoffman, 2018]. However, since our method is focused on the modeling of biological sequences, our experiments are limited to sequence data only.

#### 3.1.1 Datasets and evaluation metric

In our experiments, we consider the datasets used by Alipanahi et al. [2015], consisting of fragment peaks in 506 different ENCODE ChIP-seq experiments. While negative sequences are originally generated by random dinucleotide shuffling, we also train our models with real negative sequences not bound by the TF, a task called motif occupancy by Zeng et al. [2016]. Both datasets have a balanced number of positive and negative samples, and we therefore measure performances by the area under the ROC curve (auROC). As noted by Karimzadeh and Hoffman [2018], even though classical, this setting may lead to overoptimistic performance: the real detection problem is more difficult as it involves a few binding sites and a huge number of non-binding sites.

#### 3.1.2 Hyperparameter tuning

We discuss here the choice of different hyperparameters used in CKN and DeepBind-based CNN models.

##### Hyperparameter tuning for CNNs

In DeepBind [Alipanahi et al., 2015], the search for hyperparameters (learning rate, momentum, initialization, weight decay, DropOut) is computationally expensive. We observe that training with the initialization mechanism proposed by Glorot and Bengio [2010] and the Adam optimization algorithm [Kingma and Ba, 2015] leads to a set of canonical hyper-parameters that perform well across datasets, and to get rid of such an expensive dataset-specific calibration step. The results we obtain in such a setting are consistent with those reported by Alipanahi et al. [2015] (and produced by their software package) and by Zeng et al. [2016] (see Supplementary Figure 12 and 13). Overall, this simplified strategy comes with great practical benefits in terms of speed.

Specifically, to choose the remaining parameters such as weight decay, we randomly select 100 datasets from DeepBind’s datasets, and we use one quarter of the training samples as validation set, on which the error is used as a proxy of the generalization error. We observe that neither DropOut [Srivastava et al., 2014], nor fully connected layers bring significant improvements, which leads to an even simpler model.

##### Hyperparameter tuning for CKNs

The hyperparameters of CKNs are also fixed across datasets, and we select them using the same methodology described above for CNNs. Specifically, this strategy is used to select the bandwidth parameter *σ* and the regularization parameter *λ* (see Supplementary Figure 2 and 3), which is then fixed for all the versions of CKN and on either the DeepBind’s or Zeng et al. [2016] datasets. For unsupervised CKN, the regularization parameter is dataset-specific and is obtained by a five-fold cross validation. To train CKN-seq, we initialize the supervised CKN-seq with the unsupervised method (which is parameter-free) and use the alternating optimization update presented in section 2.2. We use the Adam algorithm [Kingma and Ba, 2015] to update the filters and the L-BFGS algorithm Zhu et al. [1997] to optimize the prediction layer. The learning rate is fixed to 0.01 for both CNN and CKN. The logistic loss is chosen to be the loss function for both this and the next protein homology detection task. All the models only use one layer. The choice of filter size, number of filters, and number of layers are also discussed in Section 3.3.

**Figure 2.**
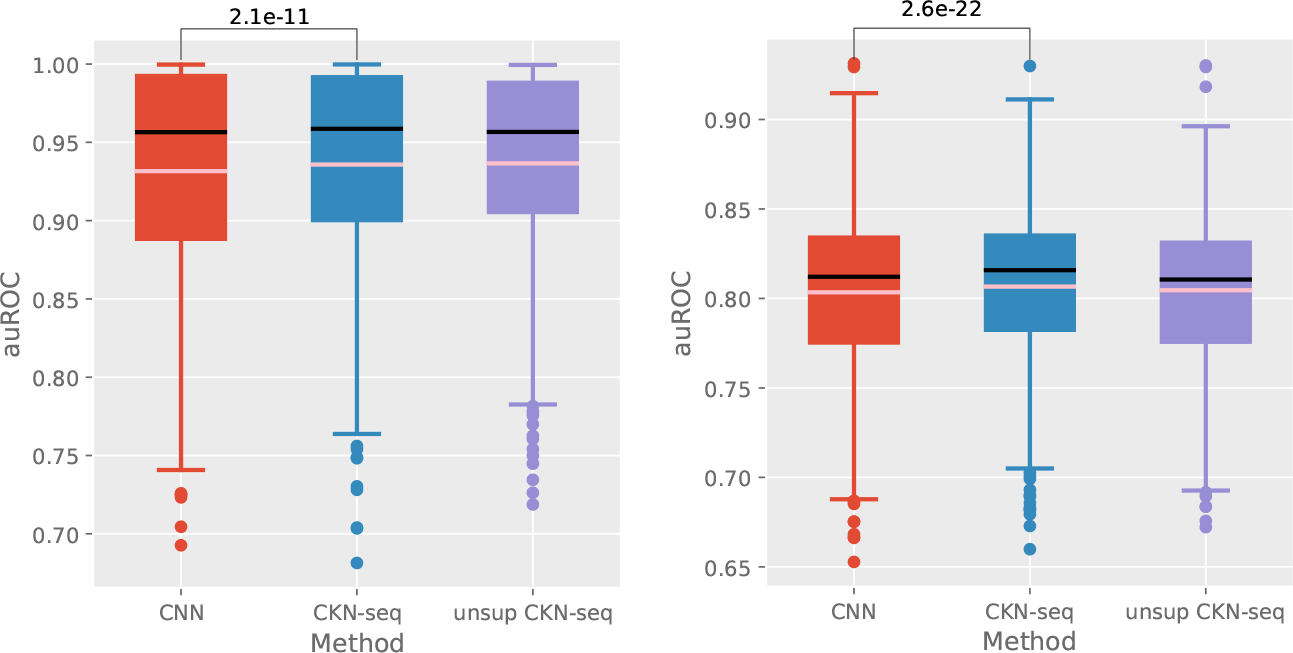
Performance comparison of CNN and CKN-seq on the DeepBind (left) and Zeng et al. [2016] (right) datasets. Number of filters for CNN and CKN-seq was set to 128 while it was 4096 for unsupervised CKN-seq. The average auROCs for CNN, CKN-seq and unsupervised CKN-seq are 0.931, 0.936, 0.937 on the DeepBind datasets and 0.803, 0.807, 0.804 on the Zeng et al. [2016] datasets. The pink and black lines respectively represent mean and median. P-values are from one-sided Wilcoxon signed-rank test. All the following figures are obtained in the same way.

**Figure 3.**
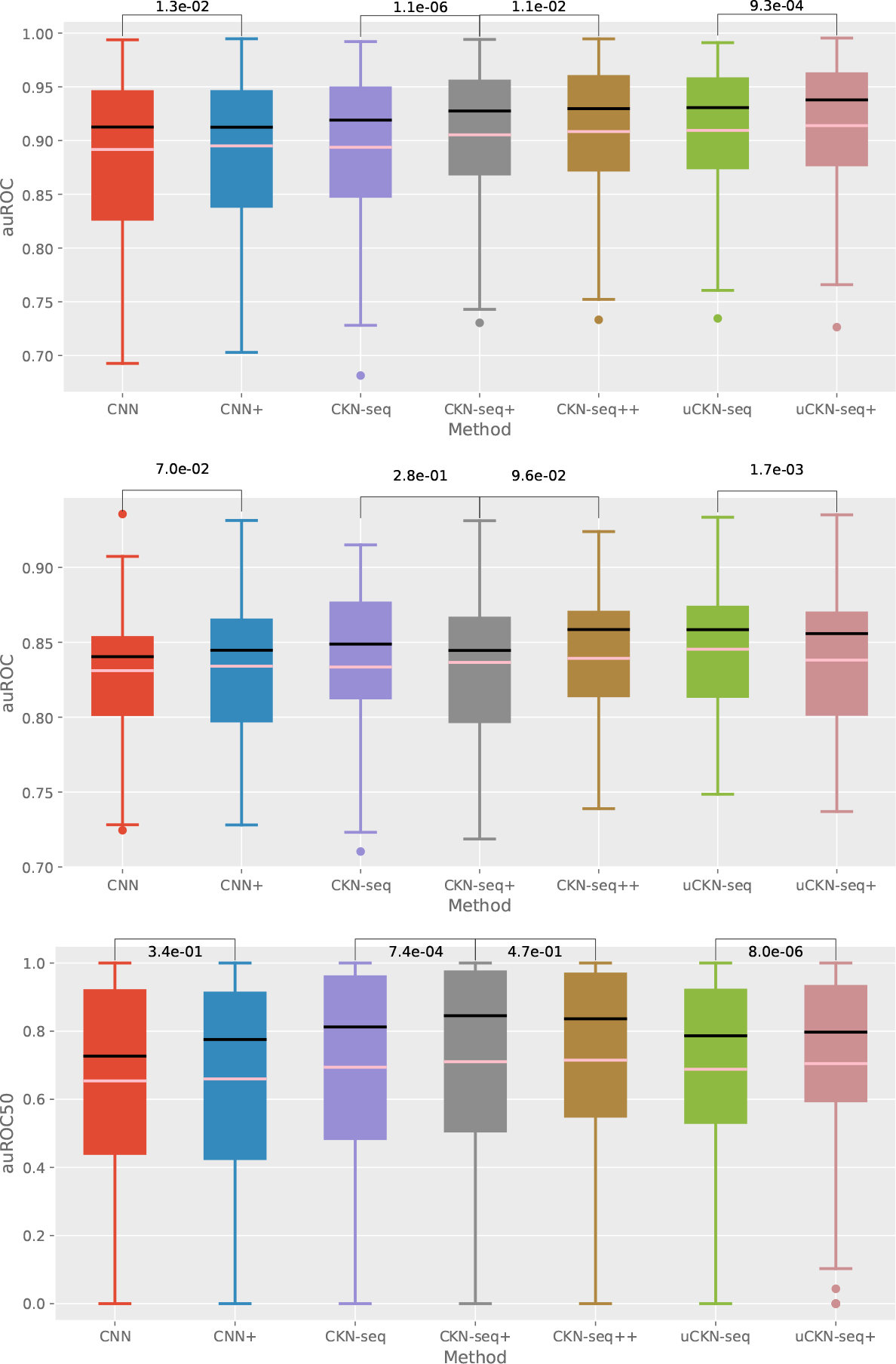
Performance comparison on small-scale datasets (top: DeepBind datasets, middle: Zeng et al. [2016] datasets, bottom: SCOP 1.67); CKN-seq+ (respectively uCKN-seq+) represents training CKN-seq (respectively uCKN-seq) with perturbation while CKN-seq++ means the hybrid model introduced in section 2.3 that combines supervised and unsupervised versions: all the models use 128 convolutional filters except that unsupervised CKN-seq (uCKN-seq) and uCKN-seq+ use 4096 filters for DeepBind’s dataset and 8192 for SCOP 1.67. The perturbation amount used in CKN-seq+, uCKN-seq+ and CKN-seq++ are respectively 0.2, 0.1 and 0.2 (0.3 for SCOP 1.67) for both tasks. The average auROC(50) for CNN+, CKN-seq++ and uCKN-seq(+) are 0.873, 0.908, 0.914 on the DeepBind datasets and 0.834, 0.839, 0.845 on the Zeng et al. [2016] datasets and 0.663, 0.715, 0.705 on SCOP 1.67.

#### 3.1.3 Performance benchmark

We compare here the auROC scores on test datasets between different CKN and DeepBind-based CNN models.

##### Performance on entire datasets

Both supervised and unsupervised versions of CKN-seq show performance similar to DeepBind-based CNN models (Figure 2), on either the Deep-Bind Zeng et al. [2016] datasets.

##### Performance on small-scale datasets

When few labeled samples are available, unsupervised CKNs achieve better predictive performance than fully supervised approaches that are hard to regularize. Specifically, we have selected all the datasets with less than 5000 training samples and reevaluated the above models. The results are presented in the top part of Figure 3. As expected, we observe that the data-augmented version outperform the corresponding unaugmented version for all the models, while the supervised CKN is still dominated by the unsupervised CKN. Finally, the hybrid version of CKN-seq presented in section 2.3 performs nearly as well as the unsupervised one while only using 32 times fewer (only 128) filters. It is also more robust to the perturbation intensity used in augmentation than the data-augmented version (detailed choice and study of perturbation intensity can be found in Supplementary Figure 4 and 5).

**Figure 4.**
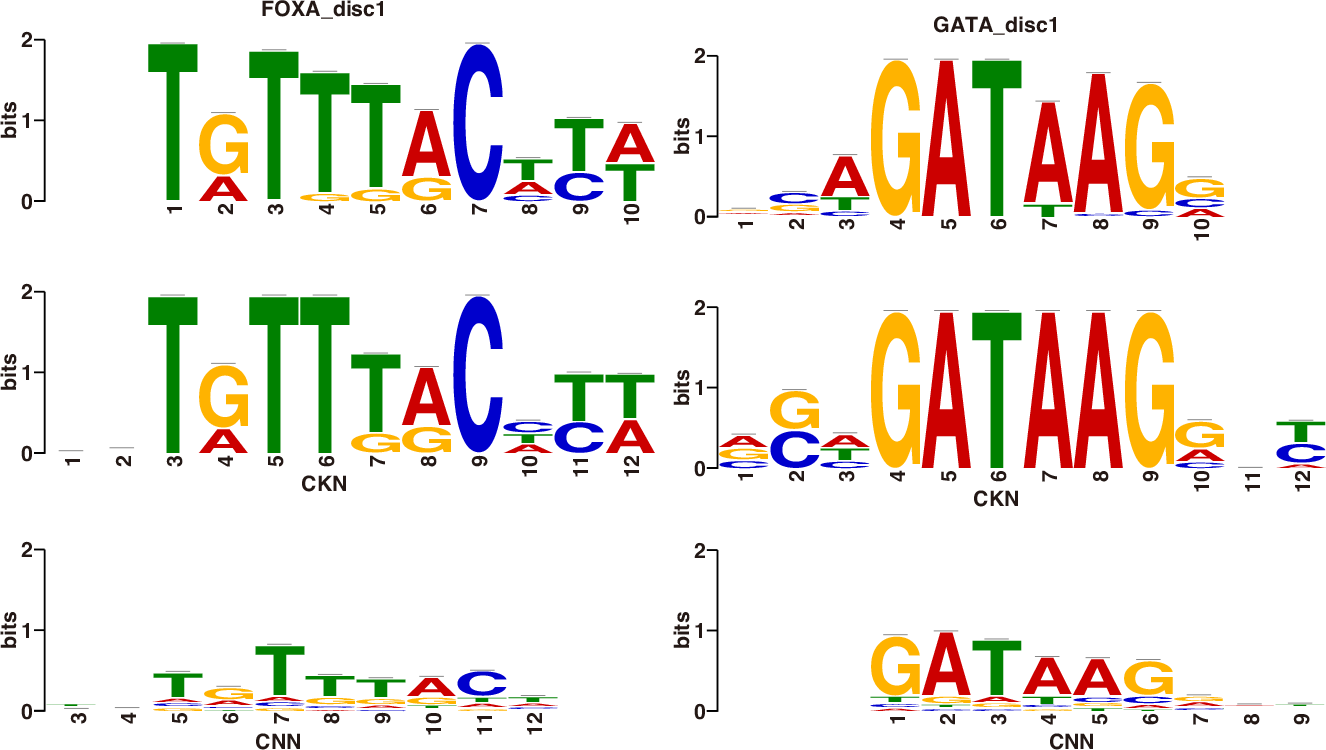
Motifs recovered by CKN-seq (middle row) and by CNN (bottom row) compared to the true motifs (top row)

We obtain similar results on the Zeng et al. [2016] datasets as shown in the middle part of Figure 3, except that the data-augmented unsupervised CKN-seq does not improve performance over its unaugmented counterpart.

### 3.2 Protein homology detection

Protein homology detection is a fundamental problem in computational biology to understand and analyze the structure and function similarity between protein sequences. String kernels, see, e.g., Leslie et al. [2002, 2004], Saigo et al. [2004], Rangwala and Karypis [2005], have shown state-of-the-art prediction performance but are computationally expensive, which restricts their use to small-scale datasets. In comparison, CKN-seq and CNN models are much more computationally efficient and also turn out to achieve better performance, which we show in the rest of this section. Specifically, we consider the remote homology detection problem and benchmark different methods on the widely-used SCOP 1.67 dataset from Hochreiter et al. [2007], including 102 superfamily recognition tasks and extending the positive training samples with Uniref50. The number of training protein samples for each task is around 5000, whose length varies from tens to thousands of amino acids. Under our formulation, positive protein sequences are taken from one superfamily from which one family is withheld to serve as test samples, while negative sequences are chosen from outside the target family’s fold.

Regarding the training of CNN and CKN-seq, we adopt the same setting as for the TF binding prediction task and the same methodology for the selection of hyper-parameters. A larger bandwidth parameter *σ* = 0.6 is selected (in contrast to *σ* = 0.3 in Section 3.1) due to the larger number of (20) characters in protein sequences. Further details about the validation scores obtained for various parameters are presented in Supplementary Figure 1-3. We also use max pooling in CKN-seq to aggregate feature vectors instead of mean pooling, which shows better performance in this problem. We fix the filter size to be 10 which seems computationally intractable for the exact algorithms, such as trie-based algorithm, for computing mismatch kernels [Kuksa et al., 2009].

Profile-based methods [Kuang et al., 2005, Rangwala and Karypis, 2005] have shown very good performance on this task but suffer a few limitations as pointed out by [Hochreiter et al., 2007], including computation time and interpretability. Nevertheless, we propose an approach which integrates profiles with CKN models. Specifically, we compute the position-specific probability matrix (PSPM) using PSI-BLAST for all the sequences in SCOP 1.67 dataset, following the same protocols as Rangwala and Karypis [2005]. PSI-BLAST is performed against Uniparc^2^ filtering out all the sequences after 2015, which leads to a database similar to the NCBI non-redundant database used by Rangwala and Karypis [2005]. We encode the sequences in Uniref50 using the BLOSUM62 position-independent probability matrix [Henikoff and Henikoff, 1992] by replacing each character with its corresponding substitution probability in BLOSUM62. Finally, we train CKN models by replacing each sequence in our kernel (2) with the square root of its corresponding PSPM (or BLOSUM62). The training and hyperparameters remain unchanged.

#### Performance on entire datasets

Besides auROC, we also use auROC50 (area under the ROC up to 50 false positives) as evaluation metric, which is extensively used in the literature [Leslie et al., 2004, Saigo et al., 2004]. Table 1 shows that unsupervised CKN-seq and CNN achieve similar performance and supervised CKN-seq achieves even better performance while they outperform all typical string kernels including local alignment kernel. They also outperform the LSTM model proposed by Hochreiter et al. [2007]. Finally, training CKN-seq is much faster than using string kernel-based methods. While training string kernel-based models requires hours or days [Hochreiter et al., 2007], training CNN or CKN-seq are done in a few minutes. In our experiments, the average training time for CNN and supervised CKN-seq is less than 3 minutes on a single cluster with a GTX1080 TI GPU and 8 CPU cores of 2.4 GHz, while training an unsupervised CKN-seq with 16384 filters (which seems to be the maximal size that can be fit to GPU memory and gives 0.956 and 0.792 respectively for auROC and auROC50) needs 30 minutes in average. We also notice that using a random sampling instead of K-means in unsupervised CKN-seq reduces the training time to 6 minutes without loss of performance. By contrast, the training time for a local alignment kernel is about 4 hours.

**Table 1.** Average auROC and auROC50 for SCOP 1.67 benchmark

| Method | auROC | auROC50 |
| --- | --- | --- |
| GPkernel [Håndstad et al., 2007] | 0.902 | 0.591 |
| SVM-pairwise [Liao and Noble, 2003] | 0.849 | 0.555 |
| Mismatch [Leslie et al., 2004] | 0.878 | 0.543 |
| LA-kernel [Saigo et al., 2004] | 0.919 | 0.686 |
| LSTM [Hochreiter et al., 2007] | 0.942 | 0.773 |
| CNN (128 filters) | 0.960 | 0.799 |
| CKN-seq (128 filters) | 0.965 | 0.819 |
| CKN-seq (128 filters) + BLOSUM62 | 0.973 | 0.835 |
| unsup CKN-seq (32768 filters) | 0.958 | 0.806 |
| Profile-based methods |  |  |
| Mismatch-profile on SCOP 1.53 [Kuang et al., 2005] | 0.980 | 0.794 |
| SW-PSSM on SCOP 1.53 [Rangwala and Karypis, 2005] | 0.982 | 0.904 |
| CKN-seq (128 filters) + profile | 0.986 | 0.906 |
| unsup CKN-seq (4096 filters) + profile | 0.968 | 0.863 |

Profile-based CKN-seq models show substantial improvements over their non-profile counterparts, including the BLOSUM62-based CKN-seq which uses the position-independent BLO-SUM62 probability matrix instead of one-hot encoding to encode sequence characters. Supervised CKN-seq shows comparable results to the best performing methods. The performance may be further improved by computing the profiles for the extended sequences in Uniref50.

#### Performance on subsampled datasets

We simulate situations where few training samples are available by subsampling only 500 class-balanced training samples for each dataset. We reevaluate the above CNN and CKN models, the data-augmented versions and also the hybrid method. The results (bottom part of Figure 3) are similar to the ones obtained for the TF binding prediction problem except that supervised version of CKN-seq performs remarkably well in this task. We also notice that CKN-seq versions trained with only 500 samples outperform the best string kernel trained with all training samples.

### 3.3 Hyperparameter Study

We now study the effect of hyperparameters and focus on the supervised version of CKN, which is more interpretable than the unsupervised one.

Both CNN and CKN-seq with one layer achieve better performance with a filter size of 12 for every fixed number of filters (Supplementary Figure 7). Since this optimal value is only slightly larger than the typical length of the motifs for TFs [Stewart et al., 2012], we deduce that the prediction mainly relies on a canonical motif while the nearby content has little contribution.

Increasing the number of filters improves the auROCs for both models regardless of the filter size, in line with the observation in Zeng et al. [2016] for CNNs. This improvement saturates when more than 128 filters are deployed, sometimes leading to overfitting (Supplementary Figure 6). We observe the same behavior for the unsupervised version of CKN-seq (Supplementary Figure 6), but usually with much larger saturation bar (larger than 4096 for TF binding prediction and 32768 for protein homology detection). When using only 16 filters, CKN-seq shows better performance than DeepBind-based CNNs. This is an advantage as large numbers of filters make the model redundant and harder to interpret.

### 3.4 Model interpretation and visualization

In this section, we study the ability of a trained CKN-seq model to capture motifs and generate accurate and informative sequence logos. We use here simulated data since the true motifs are generally not known in practice. To simulate sequences containing some given motifs represented by a PWM, we follow the methodology adopted by Shrikumar et al. [2017a] and generate 500 training and 100 test samples. We train a 1-layer CKN-seq and CNN on two tasks of the respective motif FOXA1 and GATA1 [Kheradpour and Kellis, 2013], using the same hyperparameter settings as previously. We fix the filter size and number of filters to 12 and 16 to avoid capturing too many redundant features. Both models achieve about 0.99 for the auROC on test set. The trained CNN is visualized by using the approach introduced by Alipanahi et al. [2015]. Specifically, all sequences from the test set are fed through the convolutional and rectification stages of the CNN, and only the *k*-mers that passed the activation threshold (which is 0 by default) were aligned to generate a PWM and the trained CKN is visualized by using the approach presented in section 2.4, *i.e.*, solving 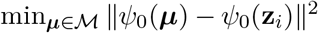 with a projected gradient descent method. The best recovered motifs (in the sense of information content) are compared to the true motifs using Tomtom [Gupta et al., 2007].

Motifs recovered by CKN-seq and CNN are both aligned to the true motifs (Figure 16). The logos given by CKN-seq are more informative and match better with the ground truth in terms of any distance measures (Table 2). This suggests that CKN-seq may be able to find more accurate motifs. We also perform the same experiments with more training samples (see Supplementary Figure 16). We observe that CKN-seq achieves small p-values in both data regimes while p-values for CNN are larger when few training samples are available.

**Table 2.** Tomtom motif p-value comparison of CKN-seq and CNN for different distance functions, see Gupta et al. [2007].

|  | FOXA1 |  | GATA1 |  |
| --- | --- | --- | --- | --- |
| Distance | CKN-seq | CNN | CKN-seq | CNN |
| KL | 8.79e-13 | 3.22e-03 | 9.94e-10 | 2.43e-03 |
| Euclidean | 1.90e-12 | 3.12e-04 | 6.25e-09 | 4.35e-04 |
| SW | 1.48e-12 | 3.83e-04 | 1.77e-09 | 4.66e-04 |
| Pearson | 1.29e-08 | 6.02e-05 | 1.37e-09 | 2.88e-04 |

## 4 Discussion

We have introduced a convolutional kernel for sequences which combines advantages of CNNs and string kernels. The resulting CKN-seq is a special case of CNN which generalizes the mismatch kernel to motifs – instead of discrete *k*-mers – and makes it *task-adaptive* and scalable.

CKN-seq retains the ability of CNNs to learn sequence representations from large datasets, leading to slightly better performance than classical CNNs on a TF binding site prediction task and on a protein homology detection task. The unsupervised version of CKN-seq keeps the kernel formalism, which makes it easier to regularize and thus leads to good performance on small-scale datasets despite the use of a huge number of convolutional filters. A hybrid version of CKN-seq performs equally well as its unsupervised version but with much fewer filters. Finally, the kernel interpretation also makes the learned model more interpretable and thus recovers more accurate motifs.

The fact that CKNs retain the ability of CNNs to learn feature spaces from large training sets of data while enjoying a RKHS structure has other uncharted applications which we would like to explore in future work. First, it will allow us to leverage the existing literature on kernels for biological sequences to define the bottom kernel *K*_0_, possibly capturing other aspects than sequence motifs. More generally, it provides a straightforward way to build models for non-vector objects such as graphs, taking as input molecules or protein structures. Finally, it paves the way for making deep networks amenable to statistical analysis, in particular to hypothesis testing. This important step would be complementary to the interpretability aspect, and necessary to make deep networks a powerful tool for molecular biology beyond prediction.

## Acknowledgements

We thank H. Zeng for sharing the experimental results of Zeng et al. [2016].

## Funding

This work has been supported by the grants from ANR (MACARON project ANR-14-CE23-0003-01 and FAST-BIG project ANR-17-CE23-0011-01) and by the ERC grant number 714381 (SOLARIS).

## Supplementary Material

In the supplementary material, we present details and additional experiments mentioned in the paper.

### A Details about the convolutional kernel

In the definition of convolutional kernel, a bottom kernel *K*_0_ was defined, for any **z**, **z′** in ℝ^*d*^, as

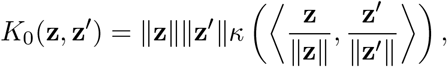

where 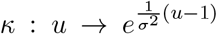 When **z** and **z′** are one-hot encoded vectors of *k*-mers, we have 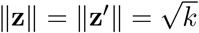 and thus

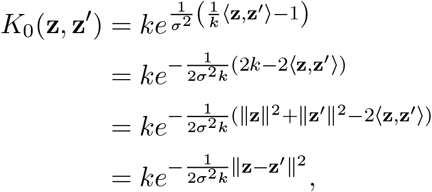

which recovers a Gaussian kernel.

### B Choice of model hyperparameters

We justify here the choice of the hyperparameters used in our experiments, including weight decay for CNNs, regularization parameter, bandwidth parameter in exponential kernel and perturbation intensity used in data-augmented CNN, CKN and hybrid model. We denote respectively by *k* the filter size and *p* the number of filters.

The scores for the following experiments are computed on a validation set, which is taken from one quarter of the training samples for each dataset and the models are trained on the rest of the training samples. For DeepBind’s datasets, we only perform validation on 100 randomly sampled datasets, which save a lot of computation time and should give similar results when using all datasets.

#### Weight decay for CNN

The choice of weight decay is validated on the validation set as shown in Figure 1.

#### Bandwidth parameter in exponential kernel

The choice of the bandwidth parameter is only validated for supervised CKN-seq and the same value is used for the unsupervised variant. Figure 2 shows the scores on the validation set when the other hyperparameters are fixed. The same choice as DeepBind’s dataset is applied to Zeng’s dataset.

#### Regularization parameter

The choice of the regularization parameter is validated following the same protocol as the bandwidth parameter. Figure 3 shows the scores on the validation set.

#### Perturbation intensity in data-augmented and hybrid model

The perturbation amount used in the data-augmented CNN, CKN and the hybrid variant of CKN are also validated on the corresponding validation set. The scores are shown in Figure 5.

**Figure 1:**
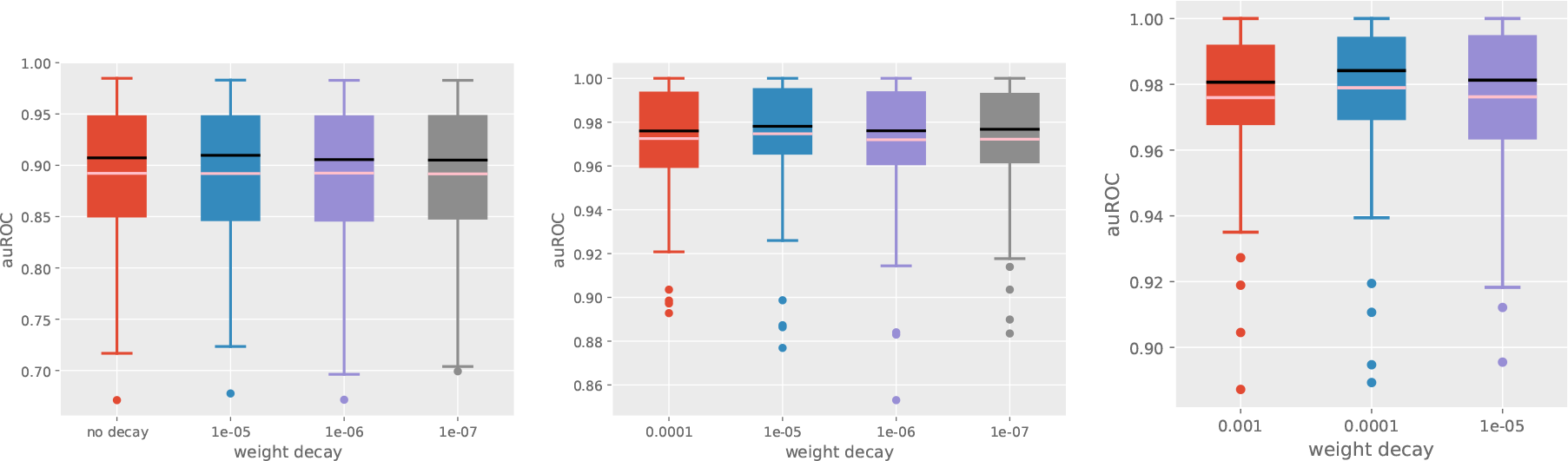
Validation of weight decay in CNNs for DeepBind’s datasets (left) and SCOP 1.67 and its subsampled datasets (middle and right); *k* = 12 and 10 respectively for each task; *p* = 128 for both tasks.

**Figure 2:**
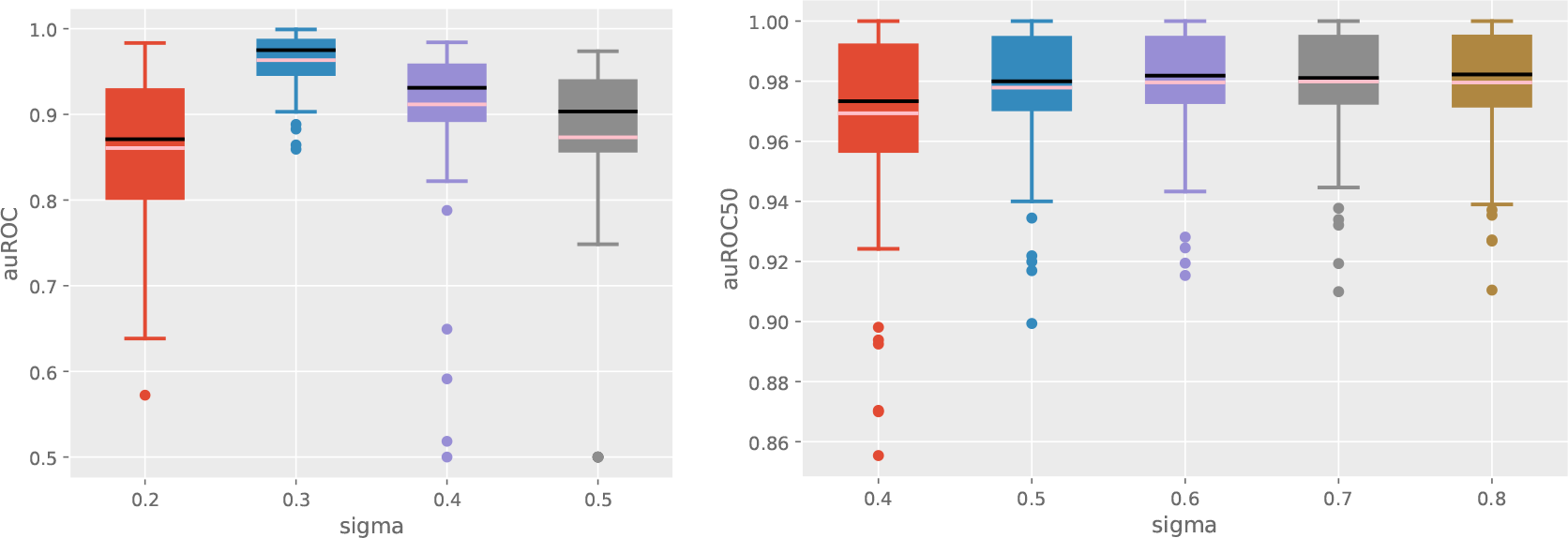
Validation of the bandwidth parameter *σ* for DeepBind’s datasets (left) and SCOP 1.67 (right). The regularization parameter is fixed to 1e-6 and 1.0 and *k* = 12 and 10 respectively for each task; *p* = 128 for both tasks.

### C Hyperparameter study

We discuss here in more detail the effect of the number and size of convolutional filters and number of layers on CNN and CKN performances. We also present the discussions on the perturbation intensity in data-augmented and hybrid variants of CKN-seq.

For some of the following comparisons, we also include the oracle model, which represents the best performance achievable by choosing the optimal parameter in comparison for each dataset (whereas parameters used in our experiments are fixed across datasets). The experiment shows that a dataset-dependent parameter calibration step could possibly improve the performance, but that the potential gain would be relatively small.

#### Number of filters, filter size and number of layers

We show in Figure 6 that increasing the number of filters improved the performance for both supervised and unsupervised variants of CKN-seq. Furthermore, the improvement of prediction performance of the supervised one was saturated when more than 128 convolutional filters were deployed.

Both CNN and CKN-seq with one layer achieve better performance with a filter size of 12 for every fixed number of filters (Figure 7). Since this optimal value is only slightly larger than the typical length of the motifs for TFs, we deduce that the prediction mainly relies on a canonical motif while the nearby content has little contribution. However if one is interested in motif discovery only, running the algorithm with larger filter size may be of interest whenever one believes that some TF binding sites are explained by larger motifs.

**Figure 3:**
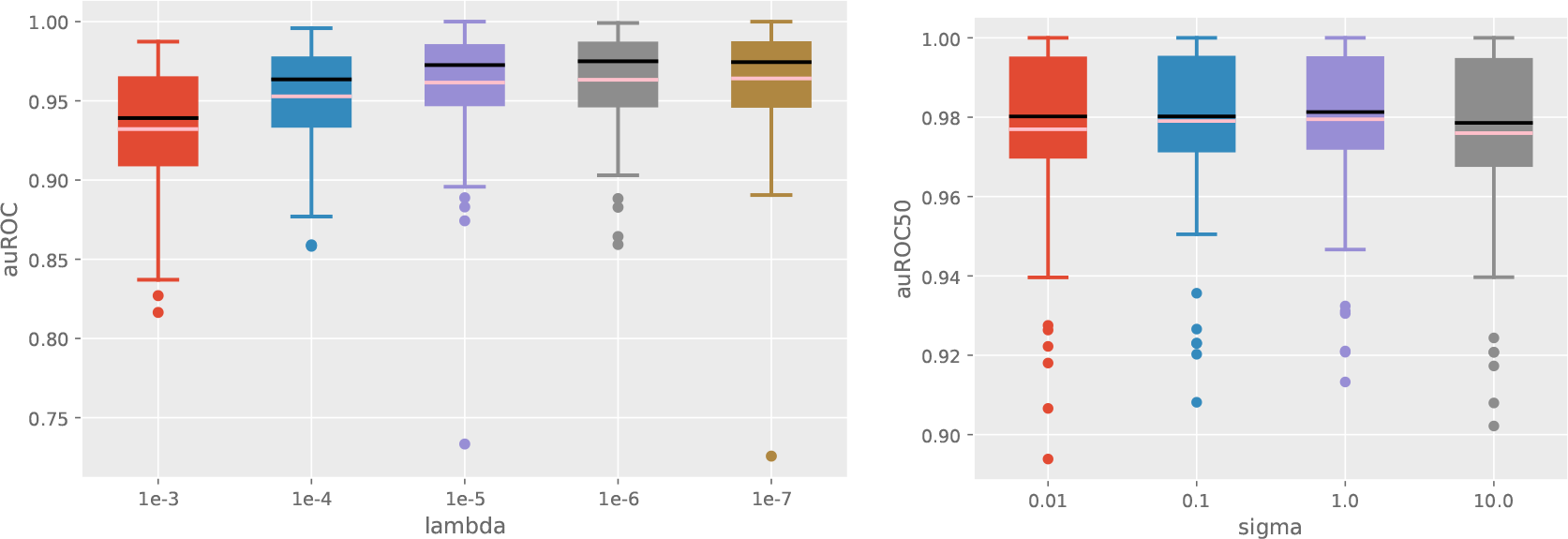
Validation of the regularization *λ* for DeepBind’s datasets (left) and SCOP 1.67 (right). The bandwith parameter is fixed to 0.3 and 0.6 and *k* = 12 and 10 respectively for each task; *p* = 128 for both tasks.

**Figure 4:**
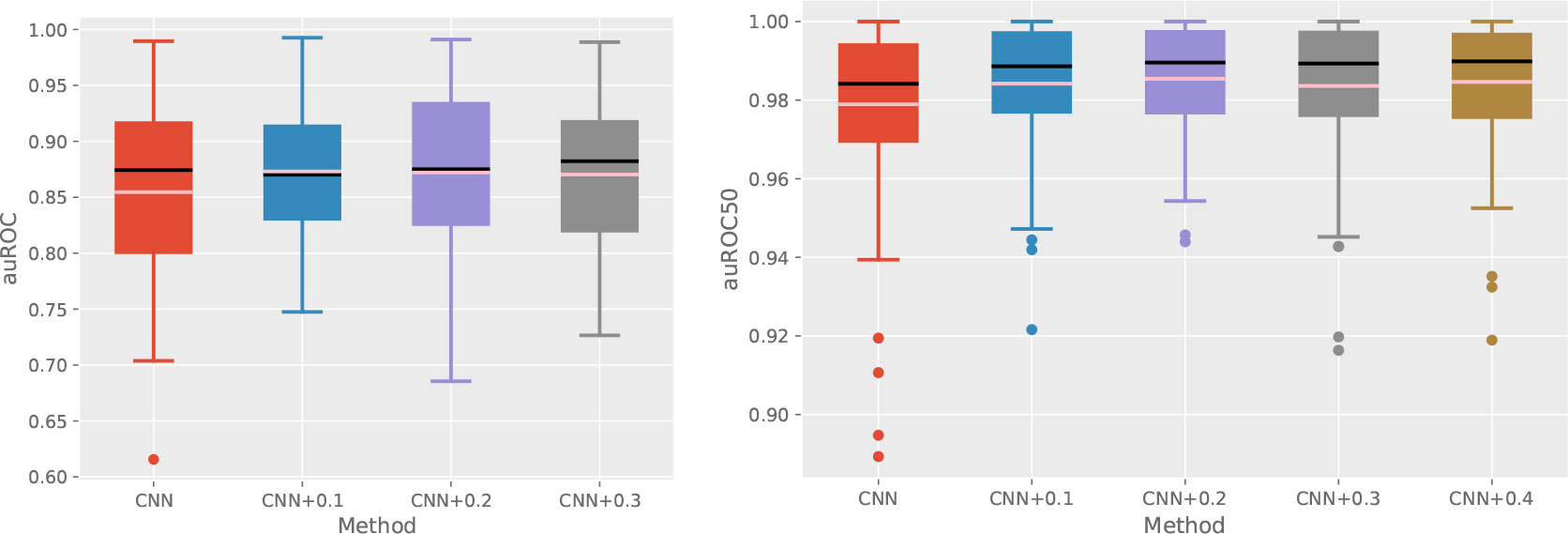
Validation of the perturbation intensity for data-augmented CNN on DeepBind’s small-scale datasets and (left) and subsampled SCOP 1.67 (right); *k* = 12 and 10 respectively for each task and *p* = 128 for both tasks.

**Figure 5:**
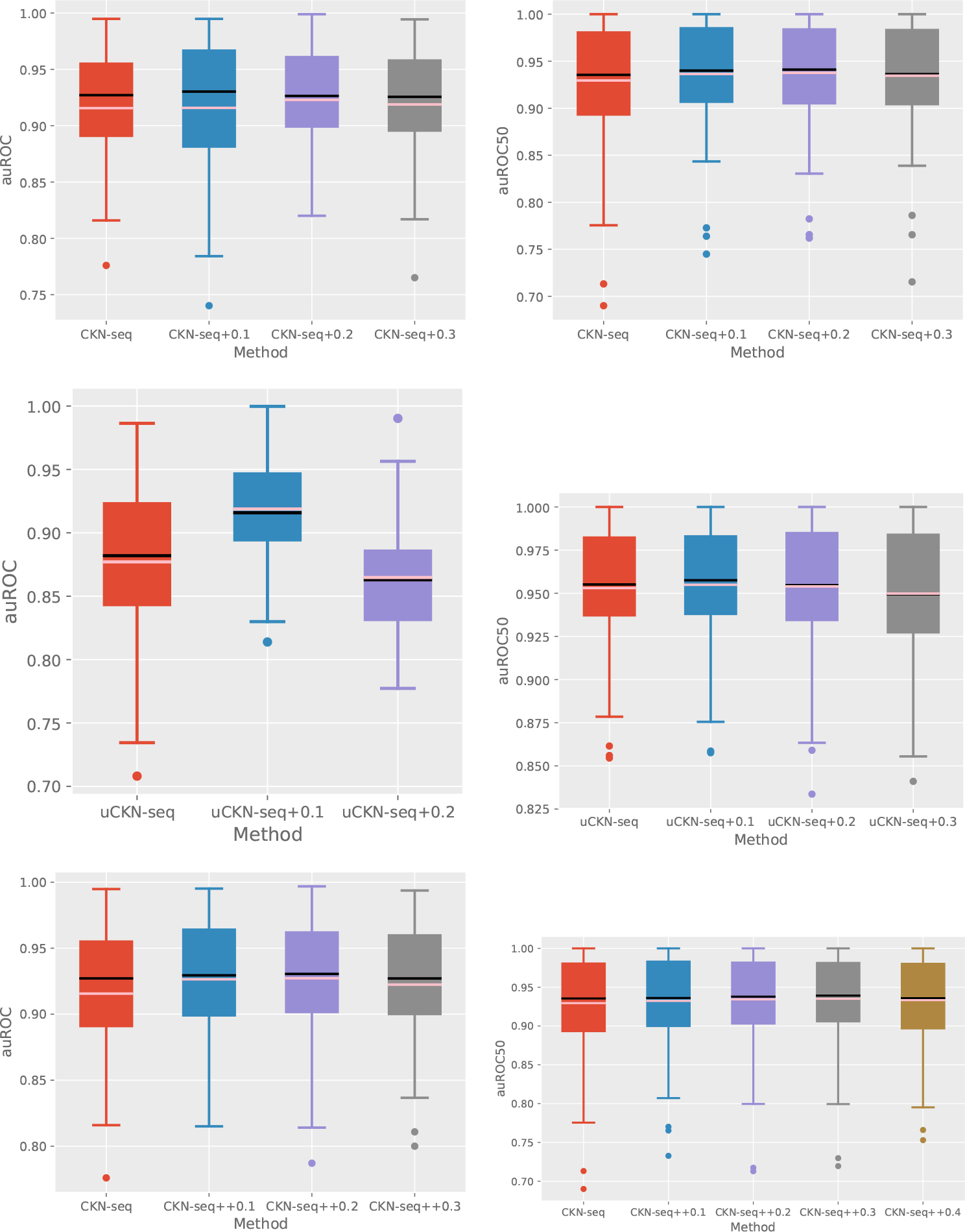
Validation of the perturbation intensity for CKN on DeepBind’s small-scale datasets (left) and subsampled SCOP 1.67 (right); each line corresponds to data-augmented supervised (top), data-augmented unsupervised (middle) and hybrid (bottom) variants of CKN-seq. The bandwith parameter is fixed to 0.3 and 0.6, the regularization parameter is fixed to 1e-6 and 1.0, and *k* = 12 and 10 respectively for each task; *p* = 128 for both tasks.

**Figure 6:**
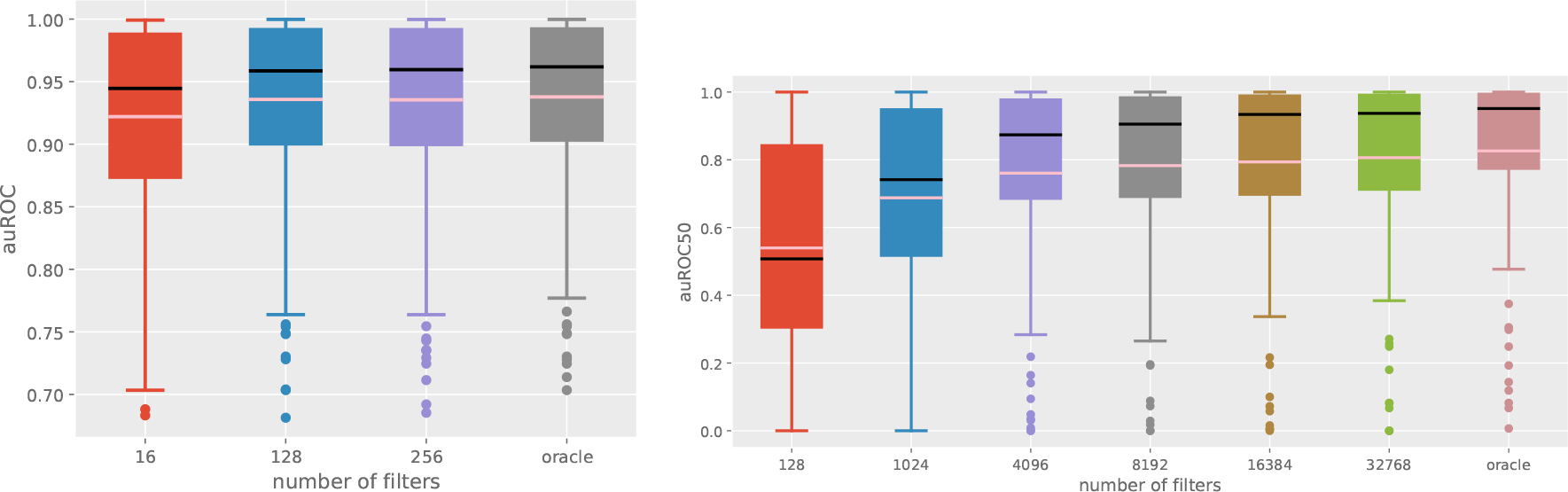
Influence of the number of filters for supervised and unsupervised CKN-seq: left supervised variant with *k* = 12 on DeepBind’s datasets; right unsupervised variant with *k* = 10 on SCOP 1.67 datasets.

**Figure 7:**
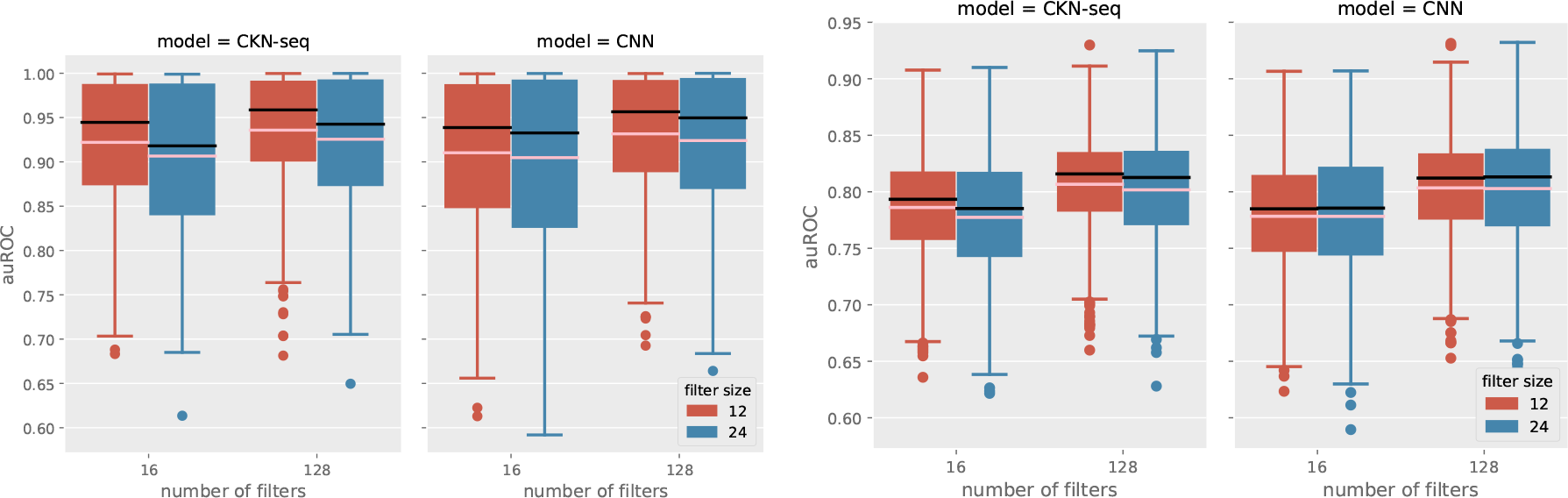
auROC scores on test datasets of DeepBind (left) and Zeng *et al.* (2016) (right) for single-layer CKN-seq and DeepBind-based CNNs with number of filters varying between 16, 64, 128 and filter size between 12, 18, 24; The pink and black line respectively represent mean and median.

Increasing the number of convolutional layers in CNNs has been shown to decrease its performance. By contrast, it does not affect the performance of CKN-seq when using a sufficient number of convolutional filters (Figure 8). Multilayer architectures allow to learn richer or more complex descriptors such as co-motifs, but may require a larger amount of data. They would also make the interpretation of the trained models more difficult. When training with 2-layer CKN models, we also notice that increasing the number of filters from 64 to 128 at the first layer or that from 16 to 64 at the second layer does not improve performance (Figure 9).

#### Perturbation intensity in data-augmented and hybrid CKN

We have shown that data augmentation improves both supervised and unsupervised CKN-seq. The hybrid approach has further improved data-augmented CKN-seq. We study here how the amount of perturbation used in augmenting training samples impacts performance. Specifically, we characterize the perturbation intensity by the percentage of changed characters in a sequence and show in Figure 11 the behavior of CKN-seq when increasing the amount of perturbation. By leveraging the best data-augmented unsupervised model on validation set, we train our hybrid variant and show its performance when increasing the amount of perturbation (Figure 10). We observe that the hybrid variant is more robust to larger amount of perturbation applied in the training samples than simply data-augmented one. Note that the results are consistent to those obtained on validation set (Section B).

**Figure 8:**
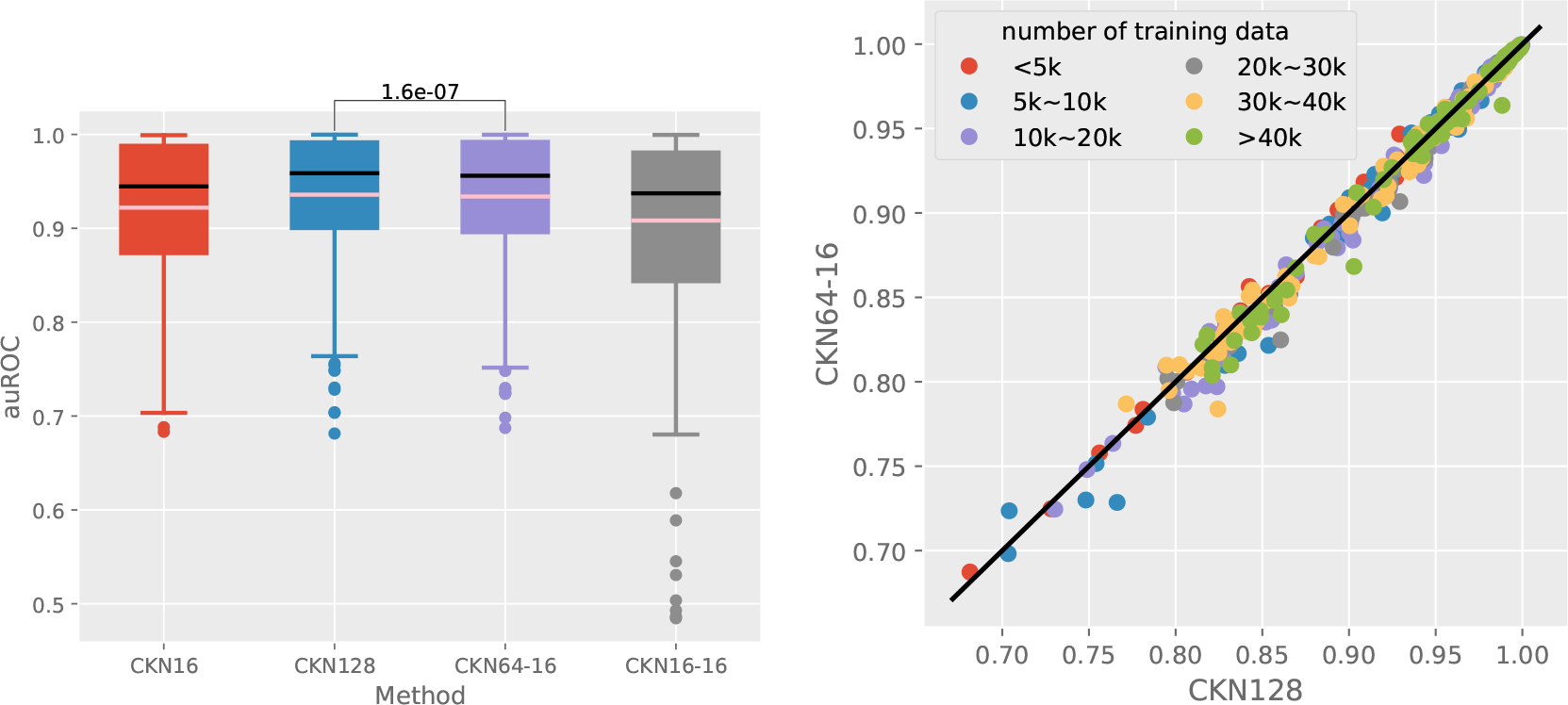
Comparison between single-layer and 2-layer CKN-seq models; note that CKN64-16 has nearly the same number of parameters as CKN128.

**Figure 9:**
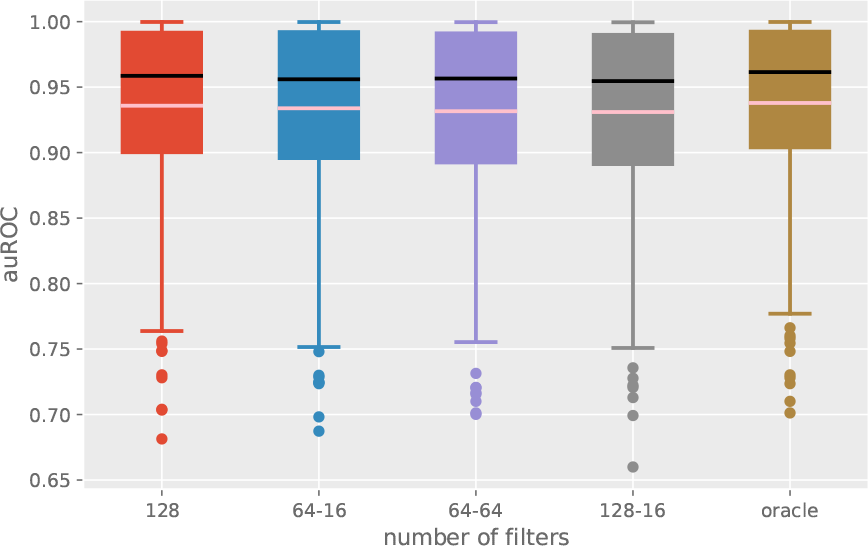
Influence of the number of filters for 2-layer supervised CKN-seq on DeepBind’s datasets.

**Figure 10:**
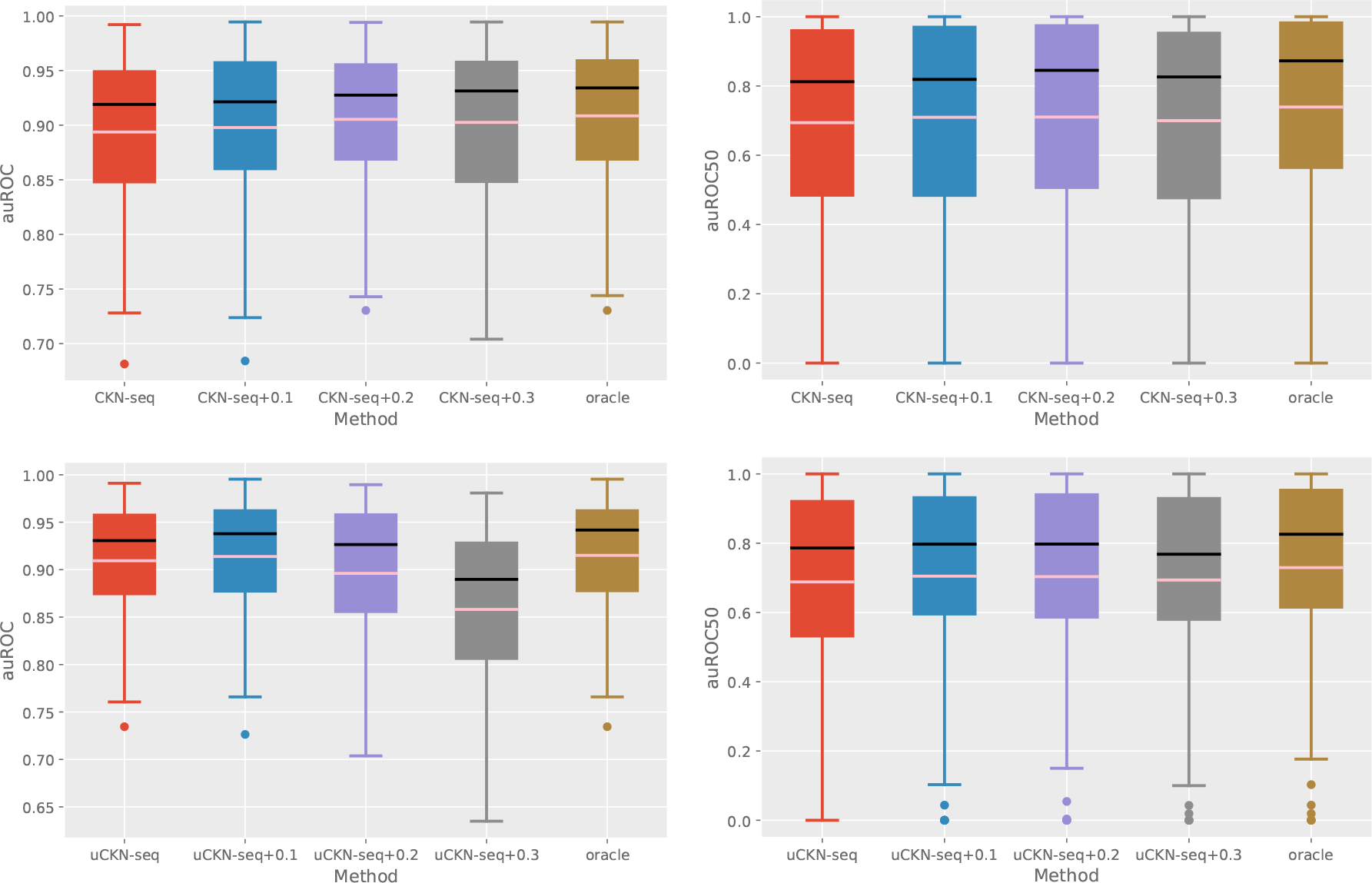
Effect of perturbation intensity on supervised and unsupervised CKN-seq: top: data-augmented supervised CKN-seq; bottom: data-augmented unsupervised CKN-seq; left: on DeepBind’s datasets; right: on SCOP 1.67. The number after + indicates the percentage of perturbation amount applied to the training samples.

**Figure 11:**
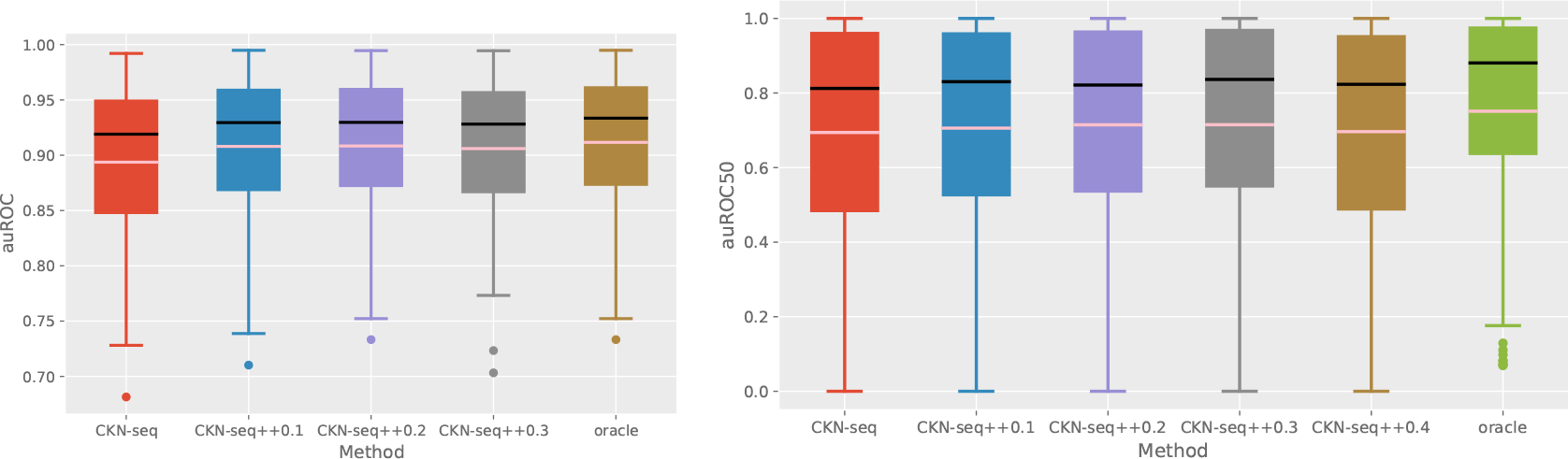
Effect of perturbation intensity on hybrid CKN-seq: left: on DeepBind’s datasets; right: on SCOP 1.67. All the hybrid models are trained using uCKN-seq+0.1. The number after ++ indicates the percentage of perturbation amount applied to the training samples.

**Figure 12:**
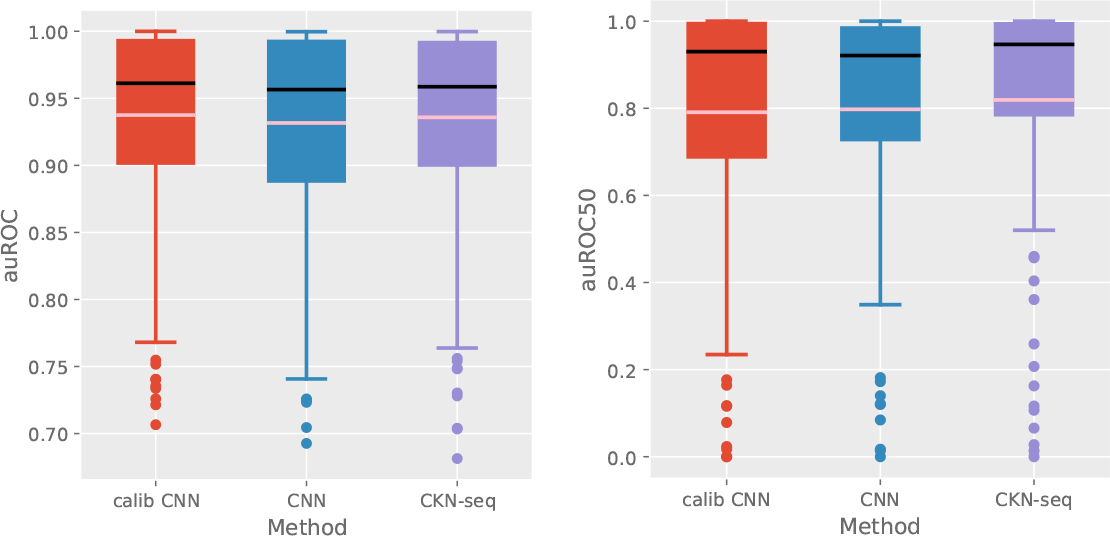
Comparison of calibrated CNN and universal models; left: DeepBind’s dataset and right: SCOP 1.67 dataset

**Figure 13:**
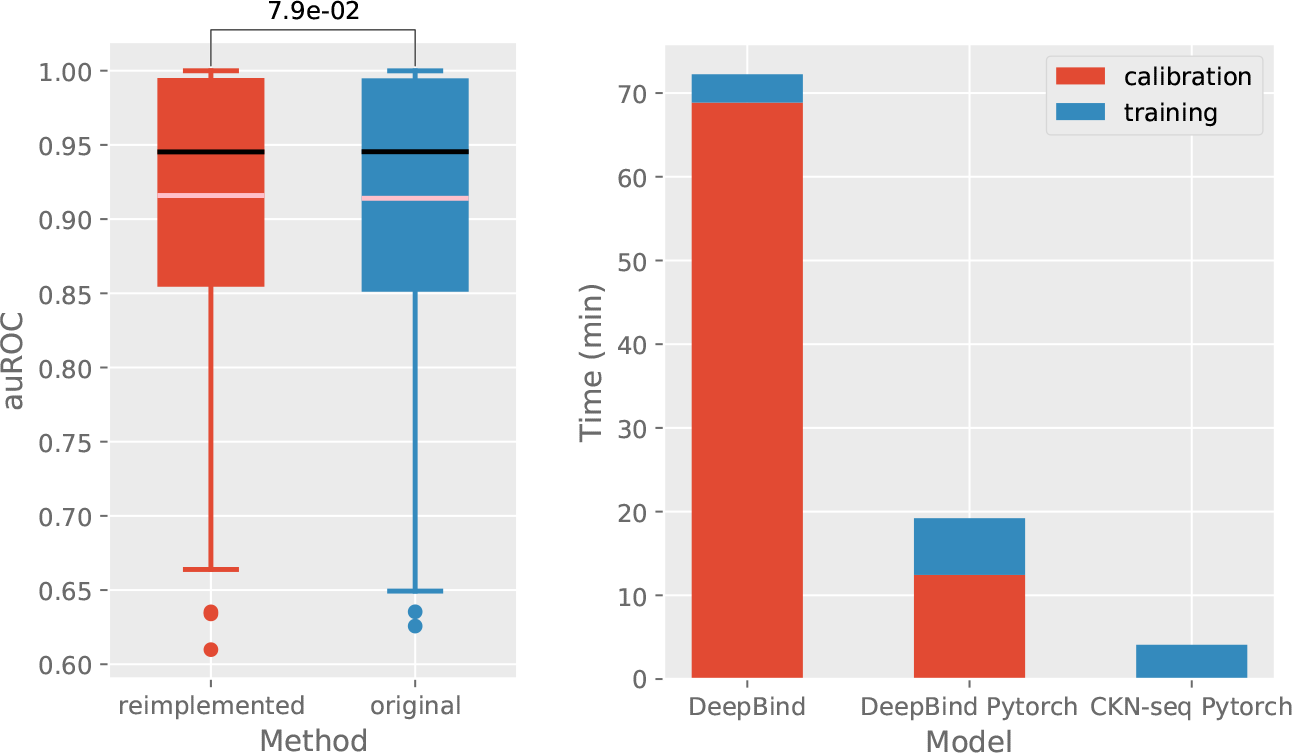
left: Comparison of reimplemented and original DeepBind, with the p-value of Wilcoxon unsigned-rank test; right: Average training time for DeepBind and CKN-seq on 50 datasets

### D Effect of hyperparameter calibration in CNN

We study here how hyperparameter calibration as used in DeepBind could affect performance and training time for CNNs. For the calibrated variant of CNN, we used the same hyperparameter search scheme used in DeepBind for the CNN, with 30 randomly chosen calibration settings and 6 training trials across the data sets.

The calibrated variant slightly outperformed hyperparameter-fixed CNN and showed similar performance to CKN-seq in the TF binding prediction task while it didn’t achieve better performance in the protein homology detection task (Figure 12).

On the other hand, training a calibrated CNN is much slower compared to hyperparameter-fixed CNN or CKN-seq. To make a fair comparison, we reimplemented and evaluated both DeepBind and CKN-seq in Pytorch. Our reimplemented model achieved almost identical performance to the original DeepBind (left panel of Figure 13) in DeepBind’s Datasets. In order to quantify the gain in training time for hyperparameter-fixed models, we measured the average training time on 50 different datasets for original DeepBind, our reimplemented DeepBind and CKN-seq on a Geforce GTX Titan Black GPU. The right panel of Figure 13 shows that training a CKN-seq model is about 25 times faster than training the original DeepBind model and 5 times faster than our reimplemented version.

**Figure 14:**
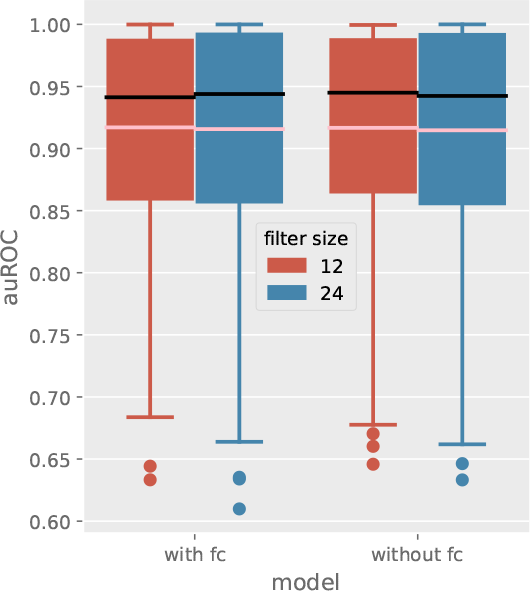
Influence of the fully connected layer in CNN on DeepBind’s datasets: all models were trained with *p* = 16.

### E Influence of fully connected layer in CNN

The authors of DeepBind have used a fully connected layer in their model. However, we found that there was no significant gain with this supplementary layer in our experiments, as shown in Figure 14.

### F Pairwise comparison of CKN and CNN

We include here some scatter plots to illustrate the pairwise comparison on each individual dataset of DeepBind and Zeng. The results are shown in 15.

### G Model interpretation and visualization

We perform the same experiments as in section 3.4 of the paper but on a larger datasets, with 9000 training samples and 1000 test samples. Motifs recovered by CKN-seq and CNN were aligned to the true motifs (Figure 16) while the logos given by CKN-seq are more informative and match better with the ground truth in terms of any distance measures (Table 3). The same conclusions can be drawn as in the small-scale case.

**Table 3.** Tomtom motif p-value comparison of CKN-seq and CNN for different distance functions.

| Distance | FOXA1 |  | GATA1 |  |
| --- | --- | --- | --- | --- |
|  | CKN-seq | CNN | CKN-seq | CNN |
| KL | 9.77e-14 | 1.78e-08 | 3.61e-11 | 3.73e-08 |
| Euclidean | 1.10e-12 | 6.62e-10 | 6.49e-12 | 1.07e-07 |
| SW | 6.75e-11 | 4.65e-10 | 1.93e-11 | 2.71e-08 |
| Pearson | 2.63e-07 | 3.59e-09 | 1.72e-08 | 5.32e-07 |

**Figure 15:**
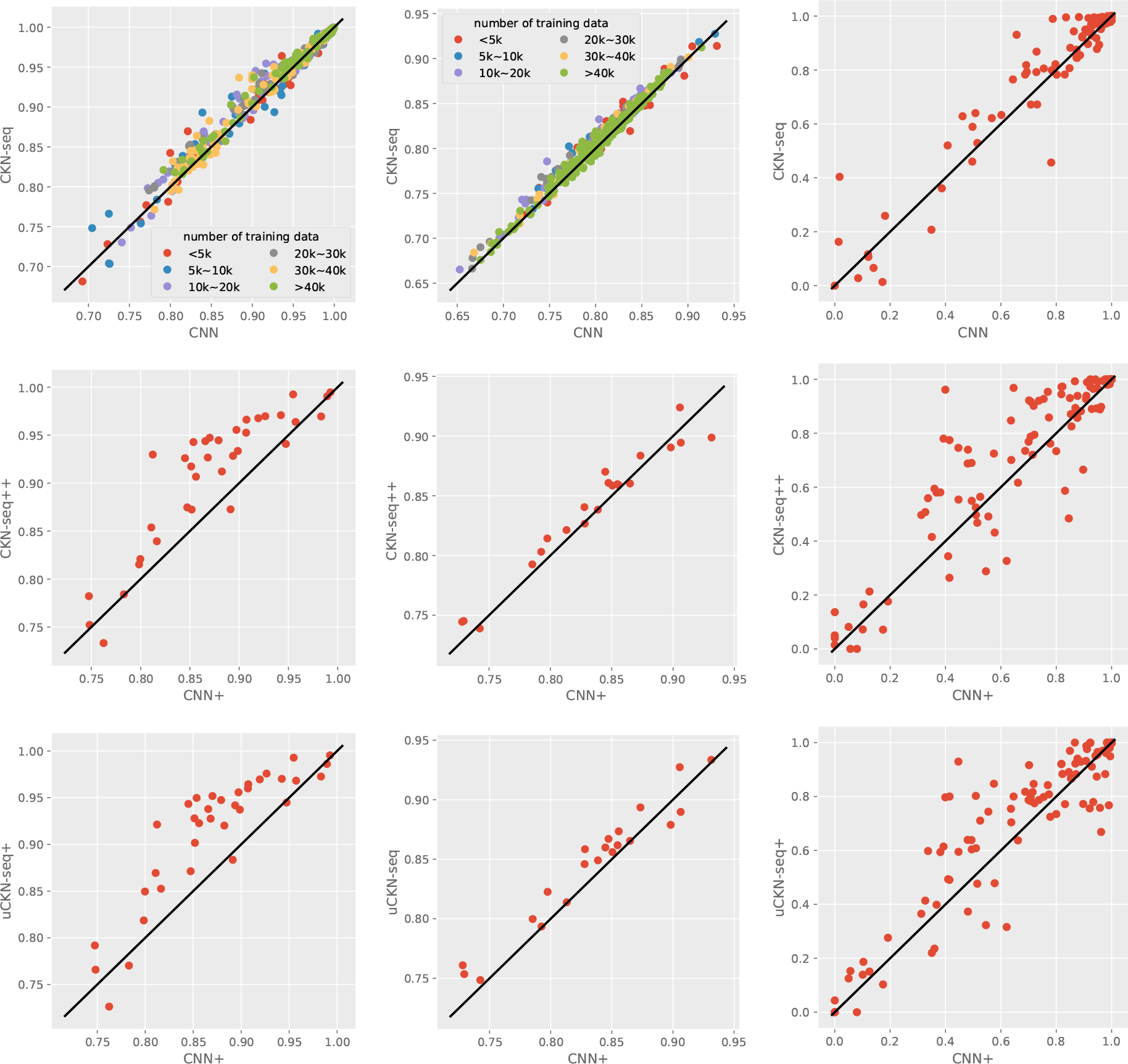
Pairwise comparison of CKN-seq and CNN on DeepBind, Zeng and SCOP 1.67 datasets. The metric is auROC for the two earlier datasets and auROC50 for the latter. The middle and bottom lines show performance of models trained on small-scale datasets.

**Figure 16:**
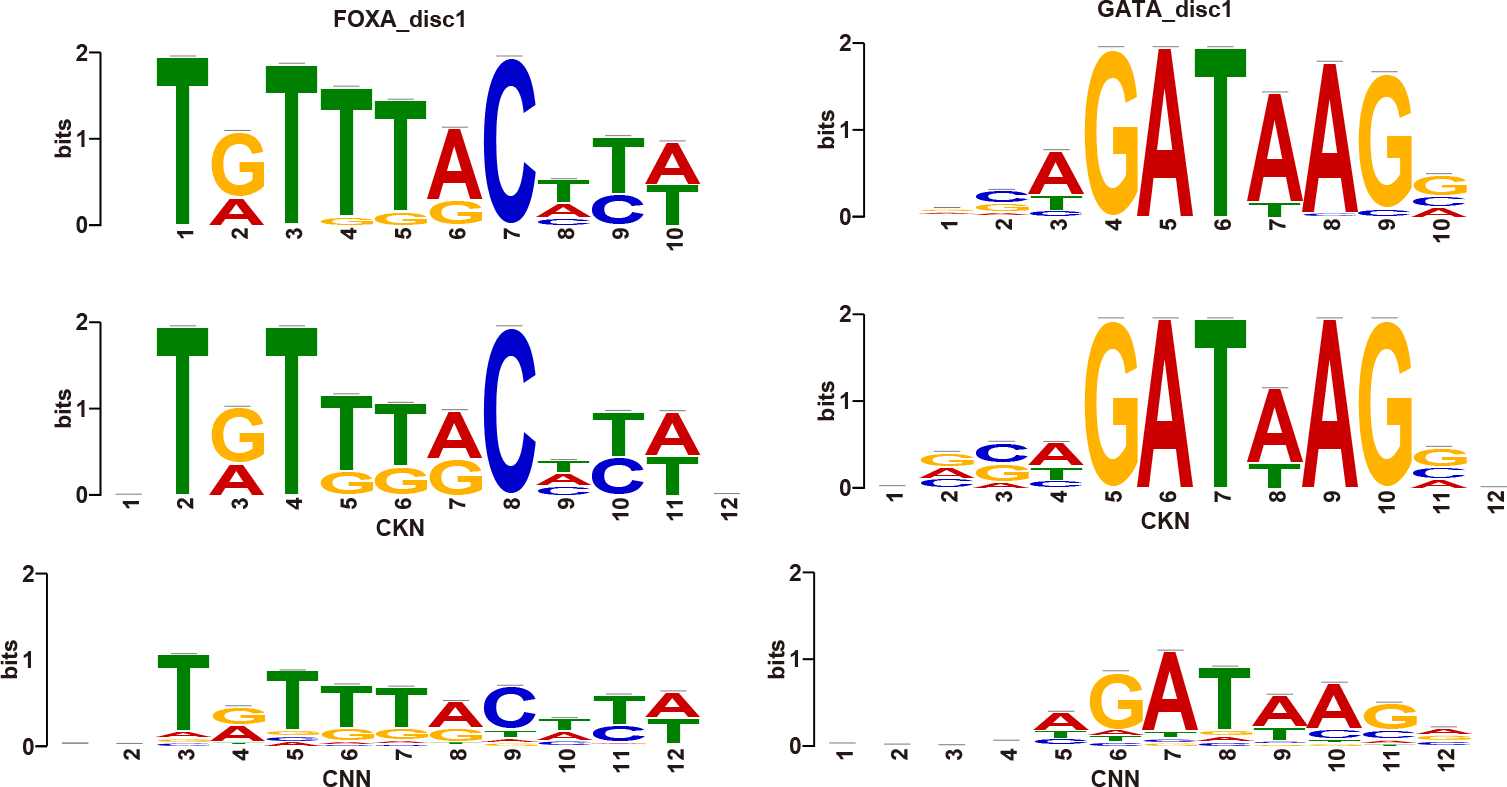
Motifs recovered by CKN-seq (middle row) and by CNN (bottom row) compared to the true motifs (top row)

### Footnotes

1 It is also possible to introduce a concept of zero-padding for sequences, such that *P*_*i*_(**x**) may contain characters outside of the original sequence, when *i* is close to the sequence boundary, see Section 3.

2 https://www.uniprot.org

## References

Babak Alipanahi, Andrew Delong, Matthew T Weirauch, and Brendan J Frey. Predicting the sequence specificities of DNA-and RNA-binding proteins by deep learning. Nature biotechnology, 33(8):831–838, 2015.

Asa Ben-Hur, Cheng Soon Ong, Sören Sonnenburg, Bernhard Schölkopf, and Gunnar Rätsch. Support vector machines and kernels for computational biology. PLoS Computational Biology, 4(10), 2008.

Alberto Bietti and Julien Mairal. Invariance and stability of deep convolutional representations. In Advances in Neural Information Processing Systems (NIPS), pages 6210–6220, 2017.

Moustapha Cisse, Piotr Bojanowski, Edouard Grave, Yann Dauphin, and Nicolas Usunier. Parseval networks: Improving robustness to adversarial examples. In International Conference on Machine Learning, 2017.

Alexandre Drouin, Sébastien Giguère, Maxime Déraspe, Mario Marchand, Michael Tyers, Vivian G Loo, Anne-Marie Bourgault, François Laviolette, and Jacques Corbeil. Predictive computational phenotyping and biomarker discovery using reference-free genome comparisons. BMC Genomics, 17(1):754, 2016.

Xavier Glorot and Yoshua Bengio. Understanding the difficulty of training deep feedforward neural networks. In Proceedings of the thirteenth international conference on artificial intelligence and statistics, pages 249–256, 2010.

Shobhit Gupta, John A Stamatoyannopoulos, Timothy L Bailey, and William Stafford Noble. Quantifying similarity between motifs. Genome biology, 8(2):R24, 2007.

Tony Håndstad, Arne JH Hestnes, and Pål Sætrom. Motif kernel generated by genetic programming improves remote homology and fold detection. BMC bioinformatics, 8(1):23, 2007.

Stephen José Hanson and Lorien Y Pratt. Comparing biases for minimal network construction with back-propagation. In Advances in Neural Information Processing Systems (NIPS), pages 177–185, 1989.

Steven Henikoff and Jorja G Henikoff. Amino acid substitution matrices from protein blocks. Proceedings of the National Academy of Sciences, 89(22):10915–10919, 1992.

Sepp Hochreiter, Martin Heusel, and Klaus Obermayer. Fast model-based protein homology detection without alignment. Bioinformatics, 23(14):1728–1736, 2007.

Tommi Jaakkola, Mark Diekhans, and David Haussler. A discriminative framework for detecting remote protein homologies. Journal of Computational Biology (JCB), 7(1-2):95–114, 2000.

Anupama Jha, Matthew R. Gazzara, and Yoseph Barash. Integrative deep models for alternative splicing. Bioinformatics, 33(14):274–282, 2017. doi: 10.1093/bioinformatics/btx268.

Mehran Karimzadeh and Michael M. Hoffman. Virtual chip-seq: Predicting transcription factor binding by learning from the transcriptome. bioRxiv, 2018. doi: 10.1101/168419. URL https://www.biorxiv.org/content/early/2018/02/28/168419.

David R Kelley, Jasper Snoek, and John L Rinn. Basset: learning the regulatory code of the accessible genome with deep convolutional neural networks. Genome Research, 26(7):990–999, 2016.

David R Kelley, Yakir Reshef, Maxwell Bileschi, David Belanger, Cory Y McLean, and Jasper Snoek. Sequential regulatory activity prediction across chromosomes with convolutional neural networks. Genome research, 2018.

Pouya Kheradpour and Manolis Kellis. Systematic discovery and characterization of regulatory motifs in encode tf binding experiments. Nucleic acids research, 42(5):2976–2987, 2013.

Diederik Kingma and Jimmy Ba. Adam: A method for stochastic optimization. 2015.

Pang Wei Koh, Emma Pierson, and Anshul Kundaje. Denoising genome-wide histone chip-seq with convolutional neural networks. Bioinformatics, 33(14):i225–i233, 2017.

Rui Kuang, Eugene Ie, Ke Wang, Kai Wang, Mahira Siddiqi, Yoav Freund, and Christina Leslie. Profile-based string kernels for remote homology detection and motif extraction. Journal of bioinformatics and computational biology, 3(03):527–550, 2005.

Pavel P Kuksa, Pai-Hsi Huang, and Vladimir Pavlovic. Scalable algorithms for string kernels with inexact matching. In Advances in neural information processing systems, pages 881–888, 2009.

Jack Lanchantin, Ritambhara Singh, Beilun Wang, and Yanjun Qi. Deep motif dashboard: Visualizing and understanding genomic sequences using deep neural networks. pages 254–265, 2017.

Yann LeCun, Bernhard Boser, John S Denker, Donnie Henderson, Richard E Howard, Wayne Hubbard, and Lawrence D Jackel. Backpropagation applied to handwritten zip code recognition. Neural computation, 1(4): 541–551, 1989.

Leslie E. Eskin, J. Weston, and W.S. Noble. Mismatch String Kernels for SVM Protein Classification. In Advances in Neural Information Processing Systems 15. MIT Press, 2003. URL http://www.cs.columbia.edu/~cleslie/papers/mismatch-short.pdf.

Christina Leslie and Rui Kuang. Fast string kernels using inexact matching for protein sequences. Journal of Machine Learning Research, 5(Nov):1435–1455, 2004.

Christina S Leslie, Eleazar Eskin, and William Stafford Noble. The spectrum kernel: A string kernel for svm protein classification. In Pacific Symposium on Biocomputing, volume 7, pages 566–575. Hawaii, USA, 2002.

Christina S Leslie, Eleazar Eskin, Adiel Cohen, Jason Weston, and William Stafford Noble. Mismatch string kernels for discriminative protein classification. Bioinformatics, 20(4):467–476, 2004.

Li Liao and William Stafford Noble. Combining pairwise sequence similarity and support vector machines for detecting remote protein evolutionary and structural relationships. Journal of computational biology, 10(6): 857–868, 2003.

C. Liu and J. Nocedal. On the limited memory bfgs method for large scale optimization. Mathematical Programming, 45(1):503–528, 1989.

Julien Mairal. End-to-end kernel learning with supervised convolutional kernel networks. In Advances in Neural Information Processing Systems (NIPS), pages 1399–1407, 2016.

Alyssa Morrow, Vaishaal Shankar, Devin Petersohn, Anthony Joseph, Benjamin Recht, and Nir Yosef. Convolutional kitchen sinks for transcription factor binding site prediction. arXiv preprint arXiv:1706.00125, 2017.

Ali Rahimi and Benjamin Recht. Random features for large-scale kernel machines. In Adv. in Neural Information Processing Systems (NIPS), pages 1177–1184, 2008.

Huzefa Rangwala and George Karypis. Profile-based direct kernels for remote homology detection and fold recognition. Bioinformatics, 21(23):4239–4247, 2005.

Hiroto Saigo, Jean-Philippe Vert, Nobuhisa Ueda, and Tatsuya Akutsu. Protein homology detection using string alignment kernels. Bioinformatics, 20(11):1682–1689, 2004.

Bernhard Schölkopf and Alexander J Smola. Learning with kernels: support vector machines, regularization, optimization, and beyond. MIT press, 2002.

Avanti Shrikumar, Peyton Greenside, and Anshul Kundaje. Learning important features through propagating activation differences. In International Conference on Machine Learning (ICML), pages 3145–3153, 2017a.

Avanti Shrikumar, Peyton Greenside, and Anshul Kundaje. Reverse-complement parameter sharing improves deep learning models for genomics. bioRxiv, 2017b.

Nitish Srivastava, Geoffrey E Hinton, Alex Krizhevsky, Ilya Sutskever, and Ruslan Salakhutdinov. Dropout: a simple way to prevent neural networks from overfitting. Journal of Machine Learning Research, 15(1): 1929–1958, 2014.

Alexander J Stewart, Sridhar Hannenhalli, and Joshua B Plotkin. Why transcription factor binding sites are ten nucleotides long. Genetics, 192(3):973–985, 2012.

Christopher KI Williams and Matthias Seeger. Using the nyström method to speed up kernel machines. In Advances in Neural Information Processing Systems (NIPS), pages 682–688, 2001.

Haoyang Zeng, Matthew D Edwards, Ge Liu, and David K Gifford. Convolutional neural network architectures for predicting DNA–protein binding. Bioinformatics, 32(12):i121–i127, 2016.

Jian Zhou and Olga Troyanskaya. Predicting effects of noncoding variants with deep learning-based sequence model. Nature Methods, 12(10):931–934, 2015.

Ciyou Zhu, Richard H Byrd, Peihuang Lu, and Jorge Nocedal. Algorithm 778: L-bfgs-b: Fortran subroutines for large-scale bound-constrained optimization. ACM Transactions on Mathematical Software (TOMS), 23(4): 550–560, 1997.

